# Increased beta2-adrenergic signaling is a targetable stimulus essential for bone healing by promoting callus neovascularization

**DOI:** 10.1101/2023.07.14.548550

**Authors:** Denise Jahn, Paul Richard Knapstein, Ellen Otto, Paul Köhli, Jan Sevecke, Frank Graef, Christine Graffmann, Melanie Fuchs, Shan Jiang, Mayla Rickert, Cordula Erdmann, Jessika Appelt, Lawik Revend, Quin Küttner, Jason Witte, Adibeh Rahmani, Georg Duda, Weixin Xie, Antonia Donat, Thorsten Schinke, Andranik Ivanov, Mireille Ngokingha Tchouto, Dieter Beule, Karl-Heinz Frosch, Anke Baranowsky, Serafeim Tsitsilonis, Johannes Keller

## Abstract

Traumatic brain injury (TBI) is associated with a hyperadrenergic state and paradoxically causes systemic bone loss while accelerating fracture healing. Here, we identify the beta2-adrenergic receptor (Adrb2) as a central mediator of these skeletal manifestations. While the negative effects of TBI on the unfractured skeleton can be explained by the established impact of Adrb2 signaling on bone formation, Adrb2 promotes neovascularization of the fracture callus under conditions of high sympathetic tone, including TBI and advanced age. Mechanistically, norepinephrine stimulates the expression of Vegfa and Cgrp primarily in periosteal cells via Adrb2, both of which synergistically promote the formation of osteogenic type-H vessels in the fracture callus. Accordingly, the beneficial effect of TBI on bone repair is abolished in mice lacking Adrb2 or Cgrp, and aged Adrb2-deficient mice without TBI develop fracture nonunions despite high bone formation in uninjured bone. Pharmacologically, the Adrb2 antagonist propranolol impairs, and the agonist formoterol promotes fracture healing in aged mice by regulating callus neovascularization. Clinically, intravenous beta-adrenergic sympathomimetics are associated with improved callus formation in trauma patients with long bone fractures. Thus, Adrb2 is a novel target for promoting bone healing, and widely used beta-blockers may cause fracture nonunion under conditions of increased sympathetic tone.

Artwork was created in BioRender.

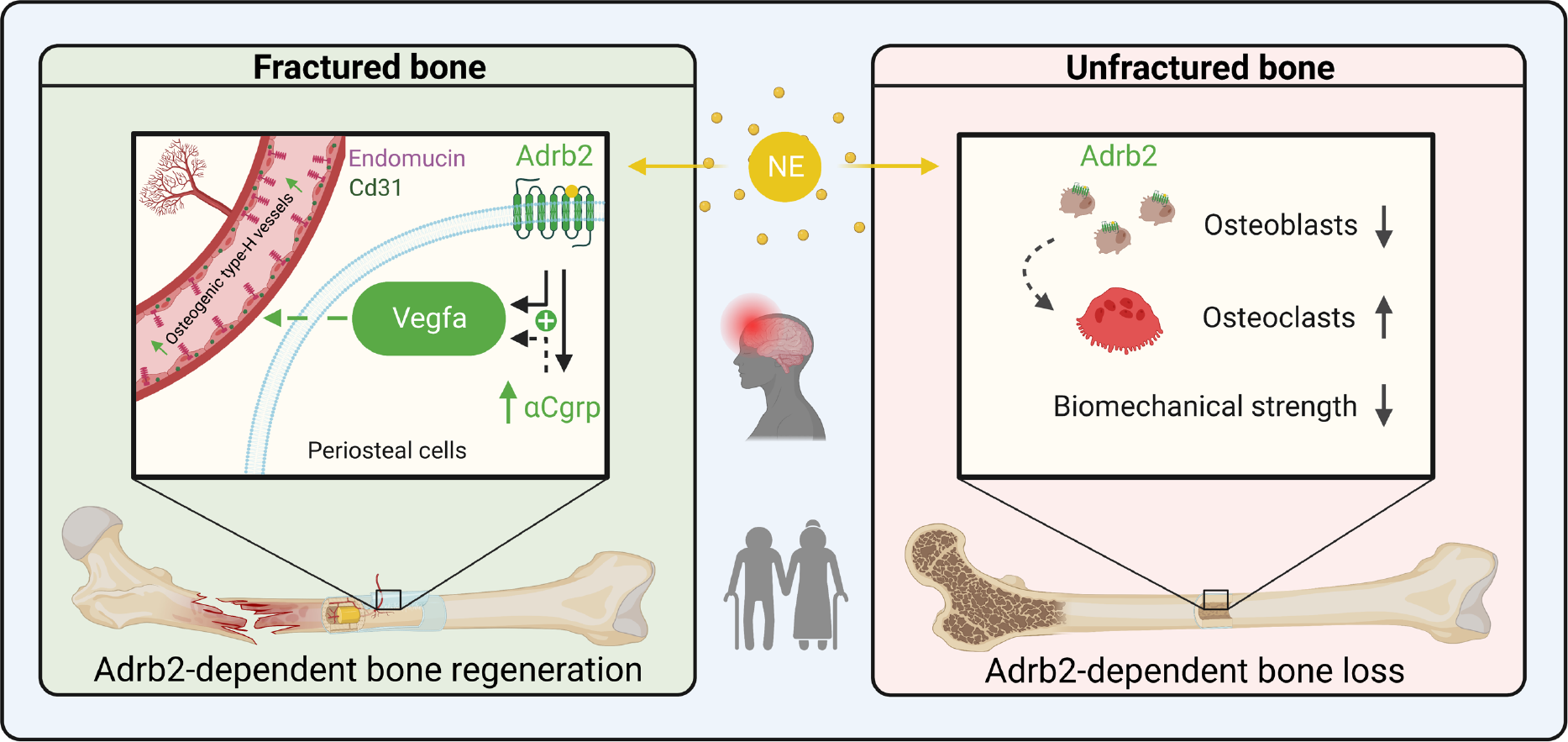

## Introduction

Bone remodeling and fracture healing are essential survival functions in young and healthy organisms that are evolutionarily conserved across a wide range of species. Bone remodeling is accomplished by the coordinated activity of osteoblasts and osteoclasts, which is regulated not only by local cytokines and growth factors, but also by systemic hormones and neurotransmitters (1). In particular, the sympathetic nervous system and its major effector molecule, norepinephrine (NE), have been identified as potent regulators of skeletal health, inhibiting bone formation and promoting bone resorption (2, 3). Similar regulatory pathways are involved in bone regeneration, allowing for scarless fracture healing in a comparatively short time. However, fracture healing is also dependent on the integrity and function of the periosteum, which circumferentially covers the bone and promotes callus neovascularization after bone injury (4).

Due to increased life expectancy, we are now faced with a high incidence of skeletal disorders such as osteoporosis and age-related fractures (5). In women, the risk of an osteoporotic fracture is equivalent to the combined risk of breast, uterine and ovarian cancer (6). Nearly 24% of hip fracture patients over the age of 50 die in the year following the fracture (6). Similarly, once a fracture occurs and is treated according to best practice, there is still a 10- 15% risk that the bone will not heal, resulting in nonunion (1). Impaired fracture healing places a heavy burden on patients, often requiring multiple revision surgeries and predisposing them to further complications, resulting in prolonged immobilization, disability, or even death (7). Many factors have been identified that contribute to impaired fracture healing. These include vascular diseases, metabolic disorders such as diabetes mellitus or osteoporosis, autoimmune disorders such as rheumatoid arthritis, immunomodulatory medications such as cortisone, and lifestyle factors such as smoking and alcohol abuse (7). In contrast, despite extensive research over the past decades, no pharmacologic agent is clinically available that reproducibly improves fracture healing and reduces the incidence of fracture nonunion with an acceptable safety profile (8).

In this context, the clinical observation that traumatic brain injury (TBI) positively affects fracture healing is of paramount importance from both a clinical and basic science perspective. In general, trauma patients with TBI have increased mortality and morbidity, including cognitive, behavioral, and physical impairments (9, 10). While TBI ultimately results in a reduction in bone density and bone quality of the unfractured skeleton (11-13), regeneration of fractured bone is paradoxically increased in these patients (14). Although several attempts have been made, a thorough pathophysiological understanding of the underlying mechanism is still lacking (13, 14). This is due to the fact that trauma to the central nervous system results in multiple and complex biological changes in the organism. For example, it is clinically known that TBI is associated with endocrine abnormalities and a hyperadrenergic state with elevated plasma catecholamine levels (15, 16). The latter is thought to reflect a generalized stress response to trauma to restore vital homeostasis in the face of TBI, as activation of the sympathetic nervous system results in massive secretion of catecholamines, including NE, to the periphery (17). In this context, it has also been shown that NE is elevated in a dose-dependent manner according to the severity of the injury, and that prolonged elevation of NE during the first 24 hours after hospital admission is highly correlated with adverse patient outcomes (18). In addition to a hyperadrenergic state, increased levels of neurotransmitters are present after TBI, some of which are known to affect bone remodeling and fracture healing including calcitonin gene-related peptide (Cgrp) (19-21). However, its role in TBI patients remained unclear.

Using our previously reported mouse model combining femoral fracture with standardized TBI (22), we show here that TBI affects both systemic bone remodeling and fracture healing through the beta2-adrenergic receptor (Adrb2). After TBI, increased NE levels reduce bone formation and promote bone resorption in the unfractured skeleton in an Adrb2-dependent manner. During bone healing, increased NE-Adrb2 signaling after TBI induces the expression of vascular endothelial factor a (Vegfa) in periosteal cells and osteoblasts, resulting in increased formation of osteogenic type-H vessels and improved vascularization of the fracture callus. While Adrb2 plays a minor role in bone healing in otherwise healthy and young mice, it significantly promotes bone regeneration in aged mice with an increased sympathetic tone. Together, our findings explain systemic bone loss and accelerated fracture healing after TBI, and identify a novel and pharmacologically targetable function of Adrb2 in fracture healing that is particularly relevant under conditions of increased sympathetic tone.

## Results

To delineate the impact of TBI on bone regeneration, we first employed our previously described mouse model combining a controlled cortical injury with a femoral osteotomy stabilized by an external fixator. Confirming our previous results (22), µCT scanning showed increased bone and tissue volume of the fracture callus of mice with TBI 14 days after surgery. (**Figure 1A**). This was confirmed on undecalcified callus sections, where an increased mineralized callus area and cartilage area were observed (**Figure 1B**). Cellular histomorphometry showed an increase in both osteoclast (**Figure 1C**) and osteoblast parameters in the callus of mice with TBI (**Figure 1D**). Assessment of the calcein-labeled area in the fracture callus demonstrated that TBI increased the formation of newly formed bone in the fracture gap (**Figure 1E**).

**Figure 1.**
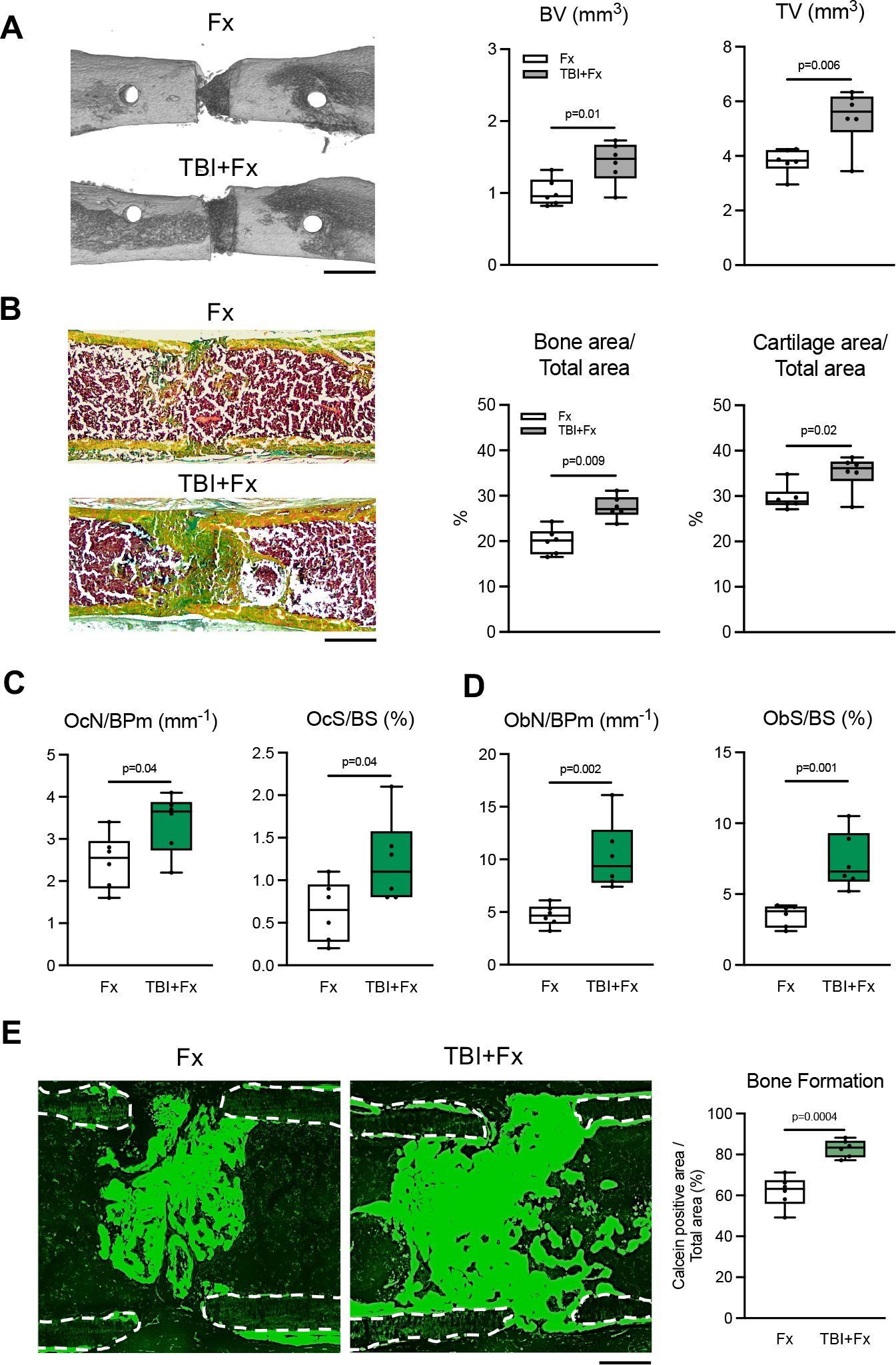
TBI improves fracture healing in mice through enhanced cellular callus remodeling. (**A**) Representative µCT images of the fractured femora 14 days following fracture only (Fx) or combined injury (TBI+Fx) in WT mice and quantification of bone volume (BV) and tissue volume (TV) in the callus. Scale bar=2 mm. (**B**) Representative callus sections (Movat Pentachrome staining) of the same mice (yellow = mineralized bone; green = cartilage) and histomorphometric quantification of bone area/total area and cartilage area/total area. Scale bar=500 µm. (**C**) Quantification of osteoclast numbers/bone perimeter (OcN/BPm) and osteoclast surface per bone surface (OcS/BS) in the same groups. (**D**) Quantification of osteoblast numbers/bone perimeter (ObN/BPm) and osteoblast surface per bone surface (ObS/BS) in the same mice. (**E**) Representative fluorescence images of calcein labeling in the fracture gap indicative of newly formed bone in the same groups. Scale bar=300µm. Quantification of calcein-positive area/total area as a marker of bone formation in the callus. For (**A**)-(**E**), data are graphed in boxplots with median and 25^th^ and 75^th^ quantiles. Whiskers indicate upper and lower extremes, respectively. n=6 female mice per group (12-week-old females, FVB/129 genetic background), two-tailed Student’s t-test.

Analysis of the unfractured bone 14 days after TBI revealed a decreased bone mass in the spine which was accompanied by a reduction in trabecular numbers and thickness (**Figure 2A**). Similar results were obtained using µCT scanning of the lumbar spine, which also demonstrated a reduced trabecular bone mass (**Supplemental Figure 1A**). Histomorphometric quantification of bone cells showed decreased osteoblast numbers and surface, whereas osteoclast parameters were increased (**Figure 2B**). In the midshaft area of the femur, cortical thickness was unaltered by TBI, however, a decreased biomechanical stability was measured using a three-point-bending test (**Supplemental Figure 1B,C**). In the distal femur, trabecular bone volume was also reduced, explained by reduced trabecular numbers and thickness (**Figure 2C**). RNA-sequencing of the unfractured midshaft femur (bone marrow not flushed out) revealed a decreased expression of osteoblast markers including *Alpl*, *Sp7*, *Ibsp*, *Bglap*, *Phex*, and *Postn* 3 days after TBI, whereas osteoclast markers were not affected (**Figure 2D**). Subsequent qRT-PCR confirmed the inhibitory impact of TBI on the expression of osteoblast markers in the unfractured femur (**Supplementary Figure 1D,E**). Together, these findings indicated that TBI primarily reduces bone formation in the unfractured skeleton while moderately increasing bone resorption. Screening RNA-seq data for a molecular target mediating the TBI effect on both bone formation and resorption, we also observed a reduced expression of the beta2-adrenergic receptor (*Adrb2*; p=0.037; adjusted p=0.550). Adrb2 was previously identified to inhibit bone formation by osteoblasts and stimulate osteoclastogenesis (23) and, as a classic example of a G-protein coupled cell surface receptor, is downregulated upon stimulation by its endogenous ligand NE (24). The downregulation of *Adrb2* in the femur of WT TBI mice was confirmed in an independent set of WT mice with TBI by qRT-PCR (**Figure 2E**). Subsequent measurement of the stable NE metabolite normetanephrine showed elevated serum levels 3 days after TBI (**Figure 2F**). Likewise, NE content in the femora of mice with TBI was increased, confirming previous reports of a hyperadrenergic state in patients with TBI in the employed model (25-30) (**Figure 2G**).

**Figure 2.**
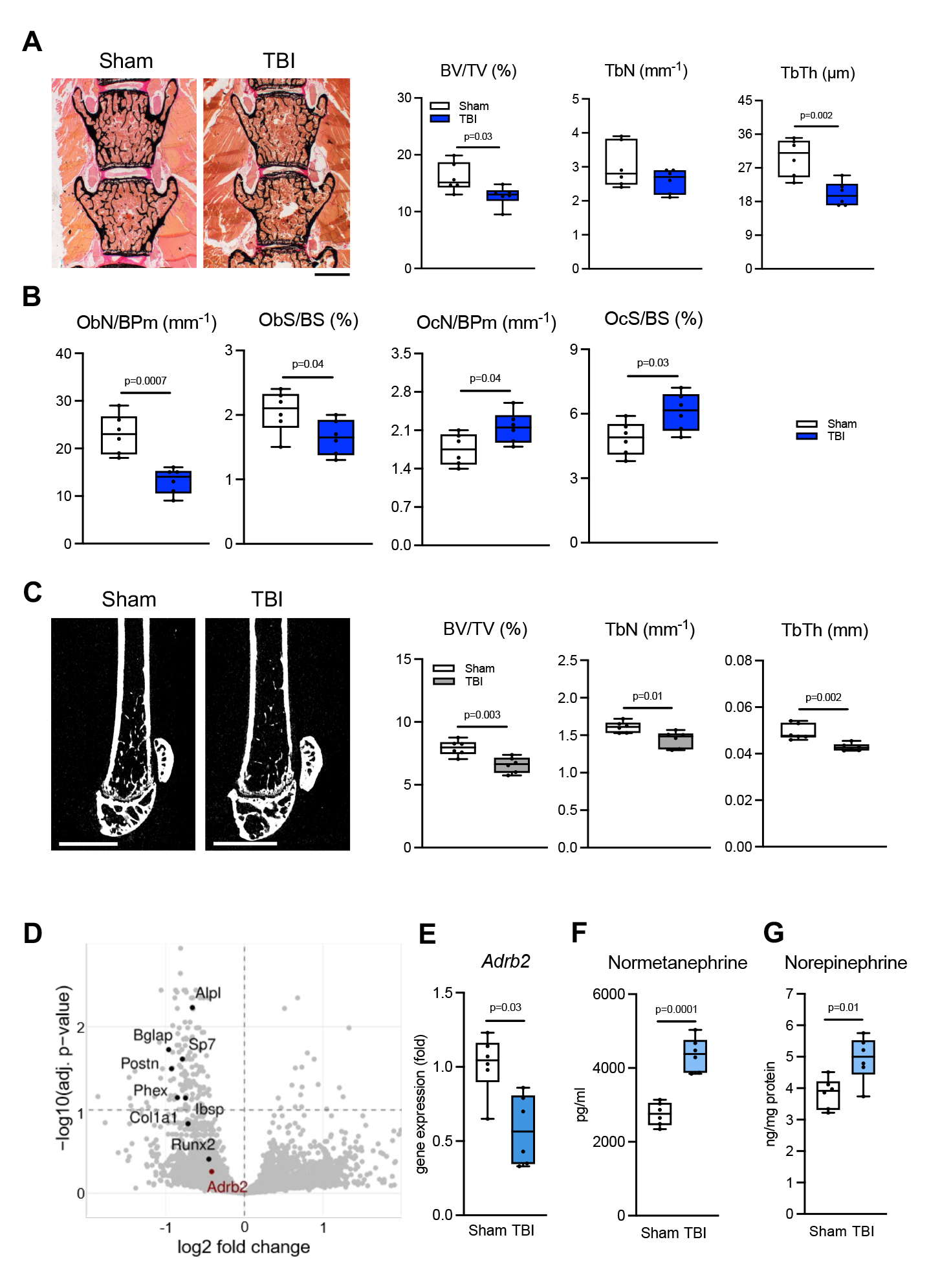
TBI causes systemic bone loss in the unfractured skeleton. (**A**) Von Kossa/van Gieson staining of undecalcified vertebra sections (L3 and L4) 14 days after sham operation or TBI and static histomorphometry of bone volume per tissue volume (BV/TV), trabecular numbers (TbN), and trabecular thickness (TbTh). n=6 mice per group (12-week-old females, FVB/129 genetic background). Scale bar=500 µm. (**B**) Quantification of osteoblast numbers/bone perimeter (ObN/BPm), osteoblast surface per bone surface (ObS/BS), osteoclast numbers/bone perimeter (OcN/BPm), and osteoclast surface per bone surface (OcS/BS). (**C**) Representative µCT images of the distal femur of the same groups and quantification of the indicated structural parameters. Scale bar=2 mm. (**D**) Summary of the RNA-seq results. The x-axis represents the relative gene expression changes between the unflushed femur midshaft of mice with sham operation (n=3 females) and TBI (n=4 females). The y-axis shows the significance level (adjusted p-value transformed in -log10 scale). (**E**) Expression of *Adrb2* in femoral midshaft samples in an independent set of WT mice with sham operation or TBI 3 days after surgery (n=6, 12-week-old females, FVB/129 genetic background). (**F**) Serum levels of normetanephrine levels 3 days following surgery in the same mice. (**G**) Norepinephrine content in the flushed femur of same groups. For (**A**)-(**C**) and (**E**)- (**G**), data are graphed in boxplots with median and 25^th^ and 75^th^ quantiles. Whiskers indicate upper and lower extremes, respectively. Two-tailed Student’s t-test.

These results indicated that the impact of TBI on the fractured and unfractured skeleton might be mediated by increased NE-Adrb2 signaling. In the distal femur of untreated WT mice, immunofluorescence with an Adrb2-specific antibody showed strong signal intensity in osteoblasts lining trabecular bone (**Figure 3A**). However, we also observed intense staining in the periosteum, which is pivotal for fracture repair. Further, in WT mice with an osteotomy, we detected an induction of *Adrb2* gene expression during the healing process, which was confirmed by immunofluorescence demonstrating increasing Adrb2 signal intensity during the fracture healing process (**Figure 3B,C**). To test the functional role of Adrb2, we next subjected 20-week-old Adrb2-deficient mice to TBI with/without a femoral osteotomy, and assessed fracture healing and bone remodeling. In the fracture skeleton, µCT-scanning demonstrated that the beneficial effect of TBI on bone healing was completely blunted in mice lacking Adrb2 (**Figure 3D**). This was confirmed in undecalcified callus sections of Adrb2-deficient mice, where no alterations in mineralized bone or cartilage area in the callus were present after TBI (**Figure 3E**). Similar observations were made in the unfractured skeleton of mice with TBI or sham operation only. In mice lacking Adrb2, bone volume and structure in the spine remained unaltered 14 days after TBI (**Figure 3F,G**). On a cellular level, osteoblast and osteoclast parameters were also unchanged in Adrb2-deficient mice with TBI compared to sham controls (**Figure 3H**). Likewise, no effect of TBI on bone architecture was found in the femora of Adrb2-deficient mice using µCT scanning (**Supplemental Figure 2A,B**). These observations were accompanied by unaltered biomechanical stability and normal expression of osteoblast and osteoclast parameters in the femoral midshaft of mutant mice (**Supplemental Figure 2C-E**).

**Figure 3.**
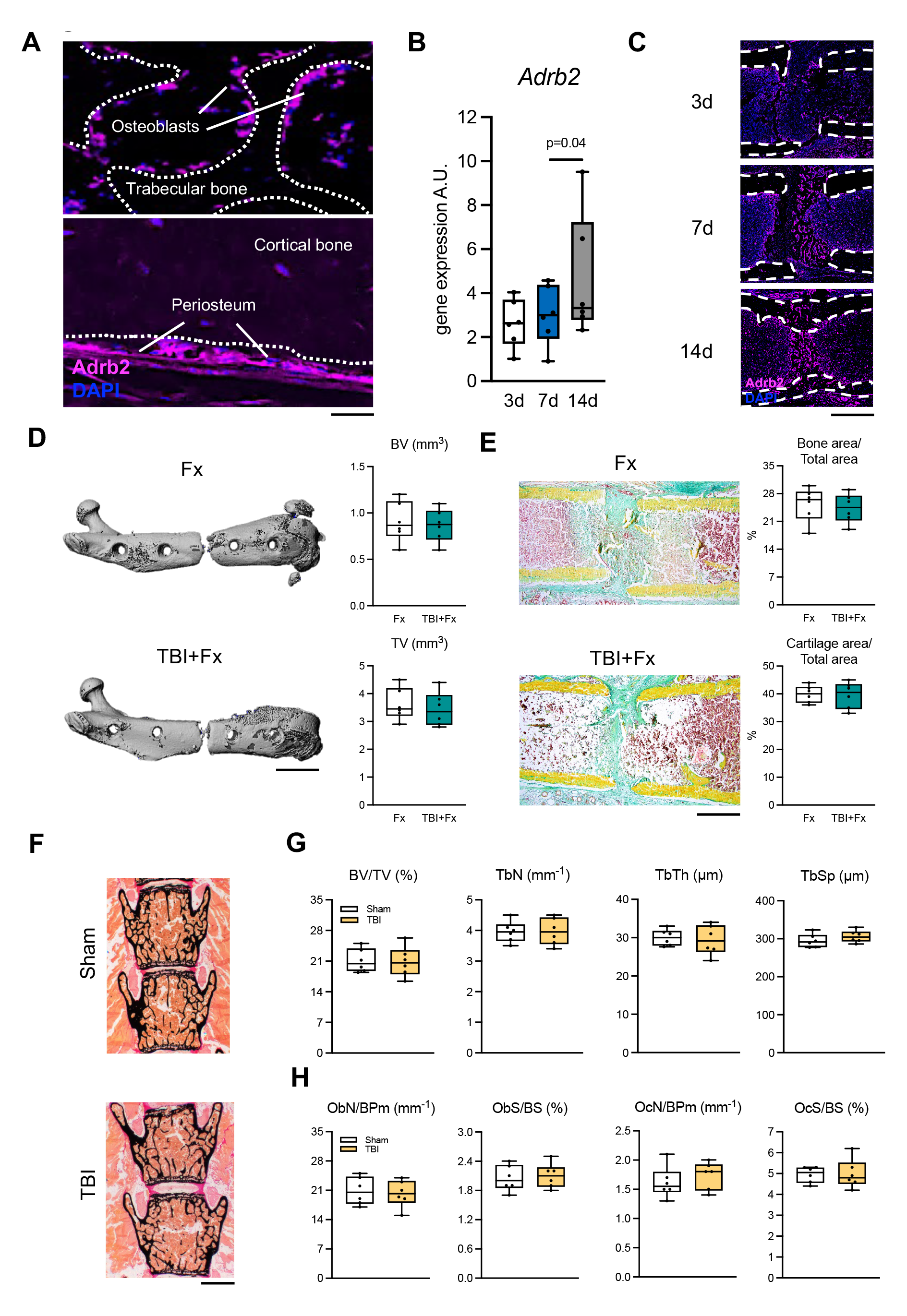
Adrb2-deficiency prevents the skeletal effects of TBI on bone remodeling and regeneration. (**A**) Representative immunofluorescent stainings using an Adrb2-specific antibody in the trabecular and cortical bone of the distal femur of 15-week-old female WT mice (C57Bl/6J genetic background). Scale bar=50 µm. (**B**) Expression of *Adrb2* in the callus at the indicated time points after osteotomy in 12-week-old WT males (n=6; FVB/129 genetic background; one-way ANOVA followed by Tukey’s post hoc test). (**C**) Adrb2-specific immunofluorescence of callus sections derived from 15-week-old WT females at the indicated time points after osteotomy (C57Bl/6J genetic background). Scale bar=400 µm. (**D**) Representative µCT images of the fractured femora 21 days following fracture only (Fx) or combined injury (TBI+Fx) in 20-week-old Adrb2-deficient females(FVB/129 genetic background) and quantification of callus bone volume (BV) and tissue volume (TV). Scale bar=3 mm. (**E**) Representative callus sections (Movat Pentachrome staining) of Fx and TBI+Fx mice 21 days after surgery (yellow = mineralized bone; green = cartilage) and histomorphometric quantification of bone area/total area and cartilage area/total area in the same mice. Scale bar=400 µm. (**F**) Von Kossa/van Gieson staining of undecalcified vertebra sections (L3 and L4) 21 days after sham operation or TBI (scale bar=500 µm), and (**G**) static histomorphometry of bone volume per tissue volume (BV/TV), trabecular numbers (TbN), and trabecular thickness (TbTh). n=6 mice per group, 20-week-old Adrb2-deficient females (FVB/129 genetic background). (**H**) Quantification of osteoblast numbers/bone perimeter (ObN/BPm), osteoblast surface per bone surface (ObS/BS), osteoclast numbers/bone perimeter (OcN/BPm), and osteoclast surface per bone surface (OcS/BS) in the same mice. All numerical data are graphed in boxplots with median and 25^th^ and 75^th^ quantiles. Whiskers indicate upper and lower extremes, respectively. For (**D**)-(**H**), two-tailed Student’s t-test was used.

Together, the above findings indicated that the hyperadrenergic state following TBI exerts opposing effects on the fractured and unfractured skeleton via Adrb2. While the inhibitory impact of Adrb2 on bone formation by osteoblasts is well-established (31), a beneficial role of Adrb2 in fracture repair was essentially unknown. We thus analyzed bone remodeling and repair in 12-week-old Adrb2-deficient mice. Undecalcified histology of the spine showed no alteration in bone architecture or cellular remodeling in Adrb2-deficient mice at this age (**Figure 4A**). Likewise, µCT-scanning of the distal femur showed no differences between mutant mice and WT littermates (**Supplemental Figure 3A**). Regarding bone healing, while showing a trend towards improved callus mineralization at an early healing stage (**Supplemental Figure 3B,C**), healing outcomes 21 days following osteotomy were not altered in Adrb2-deficient mice (**Figure 4B-D**). In sharp contrast, mutant mice at the age of 30 weeks showed an increased trabecular bone mass in the spine, which was explained by elevated indices of bone formation and reduced osteoclast parameters (**Figure 4E**). Similar observations were made in the distal femur which displayed increased bone volume (**Supplemental Figure 4A**). Assessing fracture at this age, Adrb2-deficient mice exhibited impaired callus mineralization and maturation 7 and 14 days following osteotomy on both radiological and histological levels (**Supplemental Figure 4B,C**). At 21 days after osteotomy, Adrb2-deficiency resulted in insufficient bone formation in the fracture callus, with excessive cartilage content and high rates of fracture nonunion (**Figure 4F-H**). As a cellular explanation, TRAP activity stainings showed a decrease in osteoclast parameters in the fracture callus of aged Adrb2-deficient mice (**Figure 4I**). Moreover, osteoblast numbers and surface were also dramatically reduced in aged mutant animals, accompanied by a reduction in the formation of new bone as evidenced by visualization of calcein incorporation into the healing bone (**Figure 4J**). In sum, the data show that, although comparably irrelevant for bone remodeling and regeneration at a young age, Adrb2-deficiency results in high bone mass and impaired fracture healing in the aged organism. As it is well-established that the sympathetic tone progressively increases with age (32), these findings were potentially explained by reduced Adrb2 activity due to low sympathetic tone at a young age. Although normetanephrine levels did not differ between young and aged WT mice, the expression of *Slc6a2*, mediating reuptake of NE into bone cells and thus controlling extracellular NE levels in the skeleton, was drastically reduced in aged mice in line with previous reports (33) (**Supplemental Figure 5A,B**). This was associated with an increased NE content in the femora of 30-week-old mice compared to young animals, indicative of enhanced adrenergic signaling in the skeleton of aged animals (**Supplemental Figure 5C**).

**Figure 4.**
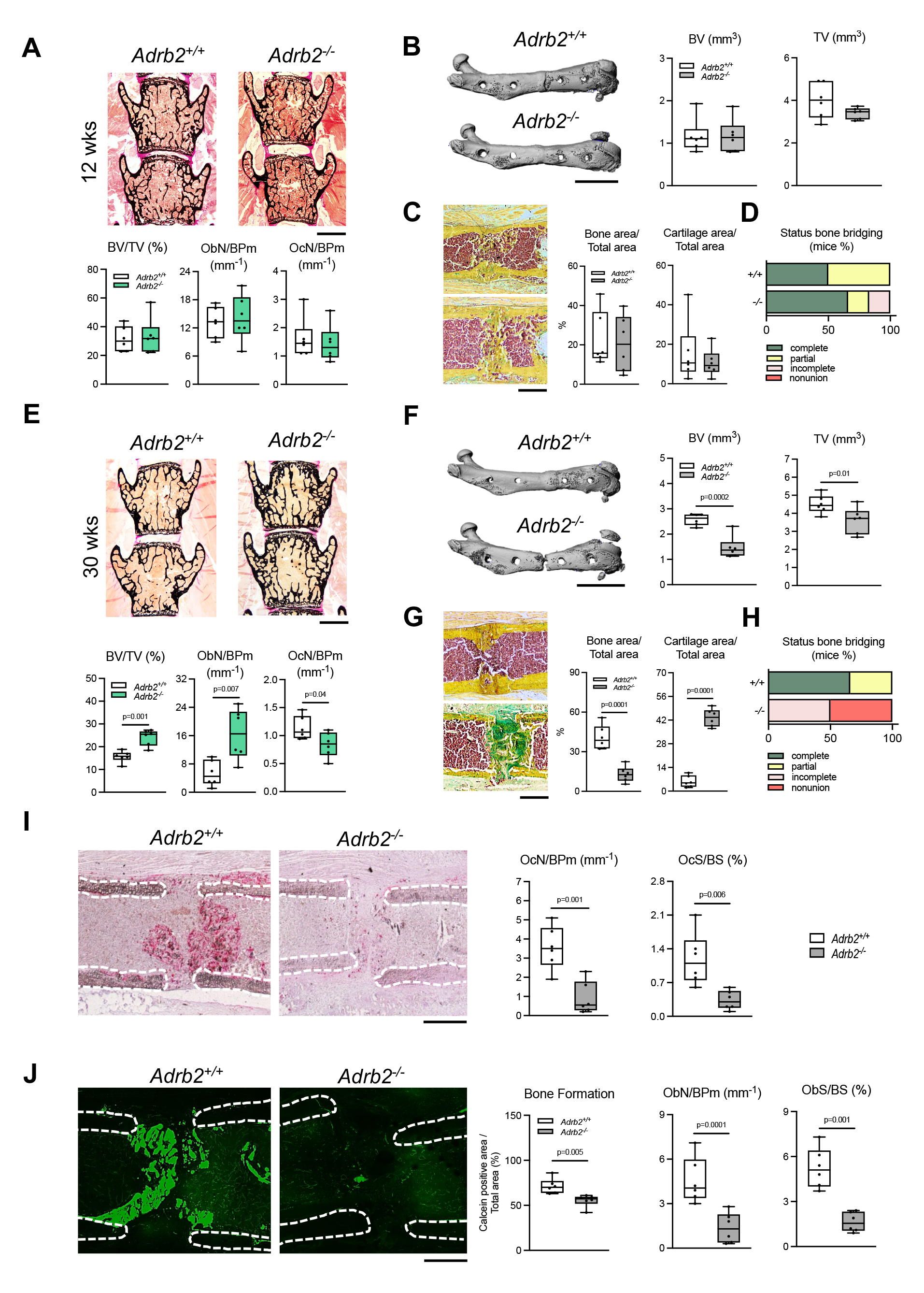
Age-dependent role of Adrb2 in fracture healing. (**A**) Von Kossa/van Gieson staining of undecalcified vertebra sections (L3 and L4) and histomorphometry of bone volume per tissue volume (BV/TV), osteoblast numbers/bone perimeter (ObN/BPm), and osteoclast numbers/bone perimeter (OcN/BPm) in 12-week-old males of the indicated genotypes. Scale bar=500 µm. (**B**) Representative µCT images of the fractured femora in 12-week-old male mice of the indicated genotypes 21 days following fracture and quantification of callus bone volume (BV) and tissue volume (TV). Scale bar=3 mm. (**C**) Representative callus sections (Movat Pentachrome staining) in the same mice and quantification of bone area/total area and cartilage area/total area. Scale bar=500 µm. (**D**) Semiquantitative evaluation of osseous callus bridging in the same mice. (**E**) Von Kossa/van Gieson staining of undecalcified vertebra sections (L3 and L4) and histomorphometry of bone volume per tissue volume (BV/TV), osteoblast numbers/bone perimeter (ObN/BPm), and osteoclast numbers/bone perimeter (OcN/BPm) in 30-week-old males of the indicated genotypes. Scale bar=500 µm. (**F**) Representative µCT images of the fractured femora in 30-week-old male mice of the indicated genotypes 21 days following fracture and quantification of callus bone volume (BV) and tissue volume (TV). Scale bar=5 mm. (**G**) Representative callus sections in the same mice and quantification of bone area/total area and cartilage area/total area. Scale bar=500 µm. (**H**) Semiquantitative evaluation of osseous callus bridging in the same mice. (**I**) Representative TRAP-stained callus sections in 30-week-old males of the indicated genotypes and quantification of OcN/BPm and OcS/BS. Scale bar=400 µm. (**J**) Representative fluorescent callus images of calcein labeling and quantification of calcein-positive area/total area in addition to ObN/BPm and ObS/BS in the same mice. Scale bar=300 µm. For (**A**)-(**J**), n=6 mice per group (FVB/129 genetic background) and two-tailed Student’s t-test were used. Except for (**D**) and (**H**), data are graphed in boxplots with median and 25^th^ and 75^th^ quantiles. Whiskers indicate upper and lower extremes, respectively.

The fact that Adrb2-deficiency blunts the beneficial impact of TBI on bone regeneration and causes fracture nonunion in otherwise healthy aged mice identifies Adrb2 as a major regulator of bone healing in conditions of increased adrenergic signaling. Because in the femur strong signals of Adrb2 were present in both osteoblasts and periosteal cells, we next monitored *Adrb2* expression in the two cell types *in vitro*. Whereas *Adrb2* was progressively induced during osteoblast differentiation, periosteal cells showed an increased expression of *Adrb2*, which declined at later stages of cell differentiation (**Figure 5A,B**). Supplementation of osteogenic culture medium with NE resulted in a decreased extracellular matrix mineralization in osteoblasts, whereas osteogenesis in periosteal cells was not affected (**Figure 5C**). In line with this, NE did not alter the expression of bone formation markers in periosteal cells, independent of whether periosteal cells were derived from calvaria or femur bones (**Figure 5D; Supplemental Figure 6A**). As the periosteum is essentially required for bone vascularization (34), a prerequisite for fracture healing, we next monitored the expression of neo-angiogenic markers. Here we found NE to robustly induce the expression of vascular endothelial growth factor a (*Vegfa*) and hypoxia-inducible factor 1-alpha (*Hif1a*) in WT but not Adrb2-deficient periosteal cells (**Figure 5E; Supplemental Figure 6B**). This effect was not only detectable on gene expression level but also translated into increased concentrations of Vegfa in the supernatant of periosteal cells (**Figure 5F**). Of note, NE also induced the expression of *Vegfa* in bone-marrow-derived osteoblast, which was however less pronounced than in periosteal cells (**Supplemental Figure 6C**).

**Figure 5.**
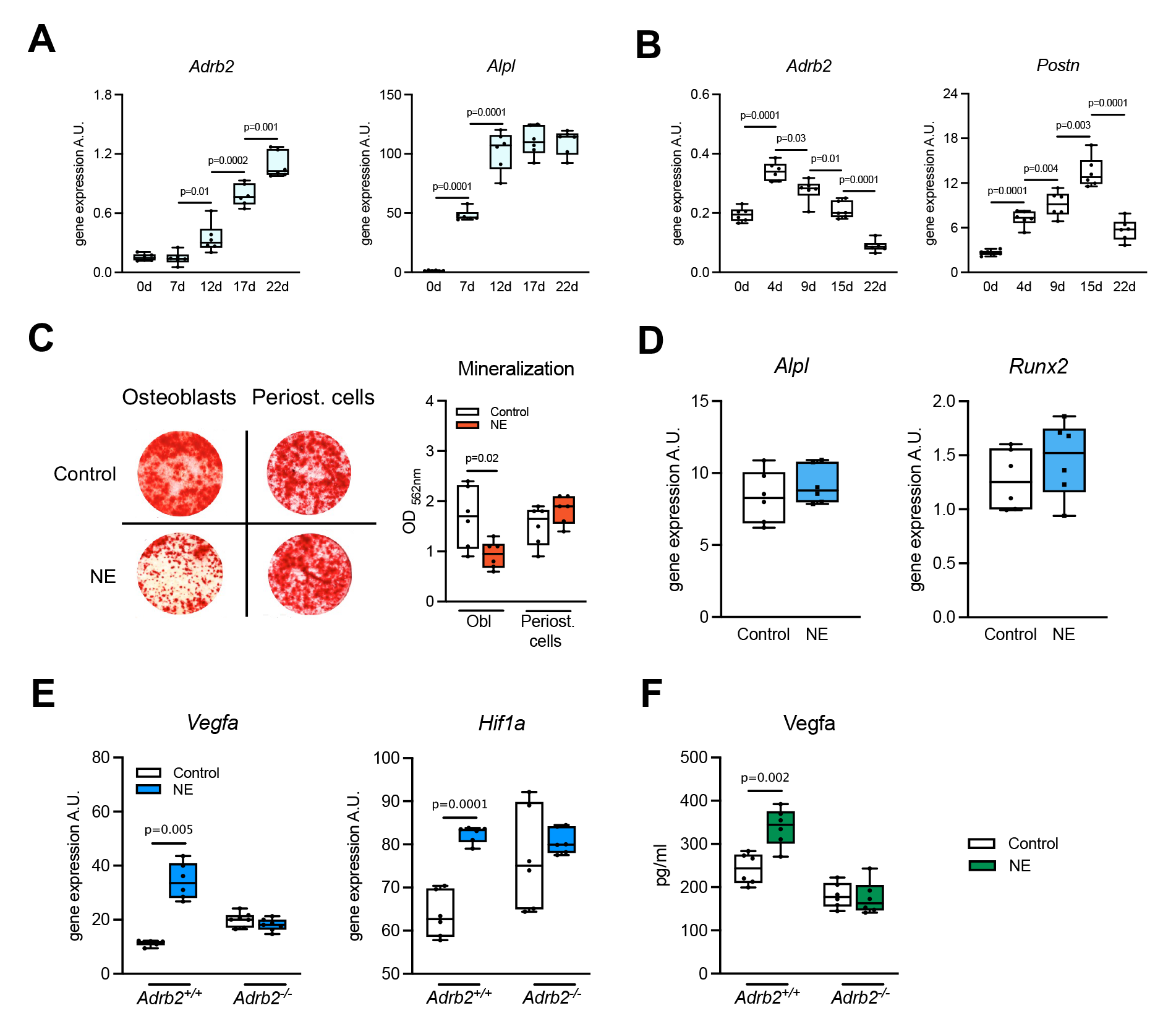
Induction of *Vegfa*-expression in periosteal cells by NE-Adrb2 signaling. (**A**) Expression of *Adrb2* and the bone formation marker alkaline phosphatase (*Alpl*) in bone marrow-derived osteoblasts and (**B**) expression of *Adrb2* and the periosteal marker periostin (*Postn*) in primary periosteal cells at the indicated time points of differentiation. One-way ANOVA followed by Tukey’s post hoc test. (**C**) Representative alizarin red stained images of bone marrow-derived osteoblasts or primary periosteal cells and quantification of extracellular matrix mineralization following 15 days of osteogenic differentiation with/without norepinephrine (NE). Two-way ANOVA followed by Tukey’s post hoc test. (**D**), (**E**) Expression of the indicated genes in WT or Adrb2-deficient periosteal cells derived from femur at day 15 of differentiation stimulated with/without NE for 6 hours. Two-tailed Student’s t-test. (**F**) ELISA of Vegfa levels in the supernatant of the same cultures. Two-way ANOVA followed by Tukey’s post hoc test. For (**A**)-(**F**), n=6 independent cultures. All cells were derived from femora of both male and female mice with FVB/129 genetic background (aged 12-18 weeks). Data are graphed in boxplots with median and 25^th^ and 75^th^ quantiles. Whiskers indicate upper and lower extremes, respectively.

To test the significance of these results *in vivo*, we next performed gene expression analysis of extracted whole callus tissue derived from aged Ardb2-deficient mice with a femoral osteotomy. Here, we observed that the expression of *Vegfa* was decreased compared to WT littermates during the very early phase of bone regeneration, i.e., 3 days after surgery, but not at day 7 (**Figure 6A**). This translated into decreased serum Vegfa levels during fracture healing at corresponding days after osteotomy (**Figure 6B**). Immunofluorescence of undecalcified cryosections of the fracture callus at day 3 confirmed a reduced Vegfa expression in the callus of mice lacking Adrb2 (**Figure 6C; Supplemental Figure 7A**). The opposite was observed in the callus of WT mice additionally subjected to TBI or sham surgery. Here, TBI resulted in an increased gene expression of *Vegfa* in the fracture callus also on day 3, but not on day 7 following surgery (**Figure 6D**), corresponding to elevated serum levels of Vegfa at the same time points (**Figure 6E**). This was again confirmed on tissue protein level using immunofluorescence with a Vegfa-specific antibody, demonstrating a higher signal intensity in the callus of TBI mice compared to sham controls (**Figure 6F; Supplemental Figure 7B)**. It is now well-established, that Vegfa is pivotal to the formation of type-H vessels, which essentially link osteogenesis to vascularization and are characterized by the co-expression of Cd31 and Endomucin in regenerating bone (35, 36). Therefore, we next studied type-H vessel formation at the end of the vascularization stage in fracture healing, corresponding to 14 days after osteotomy in mice. Here, Adrb2-deficient mice showed more than a 50% reduction in the formation of type-H vessels in the callus at this time point (**Figure 6G; Supplemental Figure 8A**). In turn, increased formation of type-H vessels was detected in the fracture callus of mice with a concomitant TBI at the same healing stage (**Figure 6H; Supplemental Figure 8B**).

**Figure 6.**
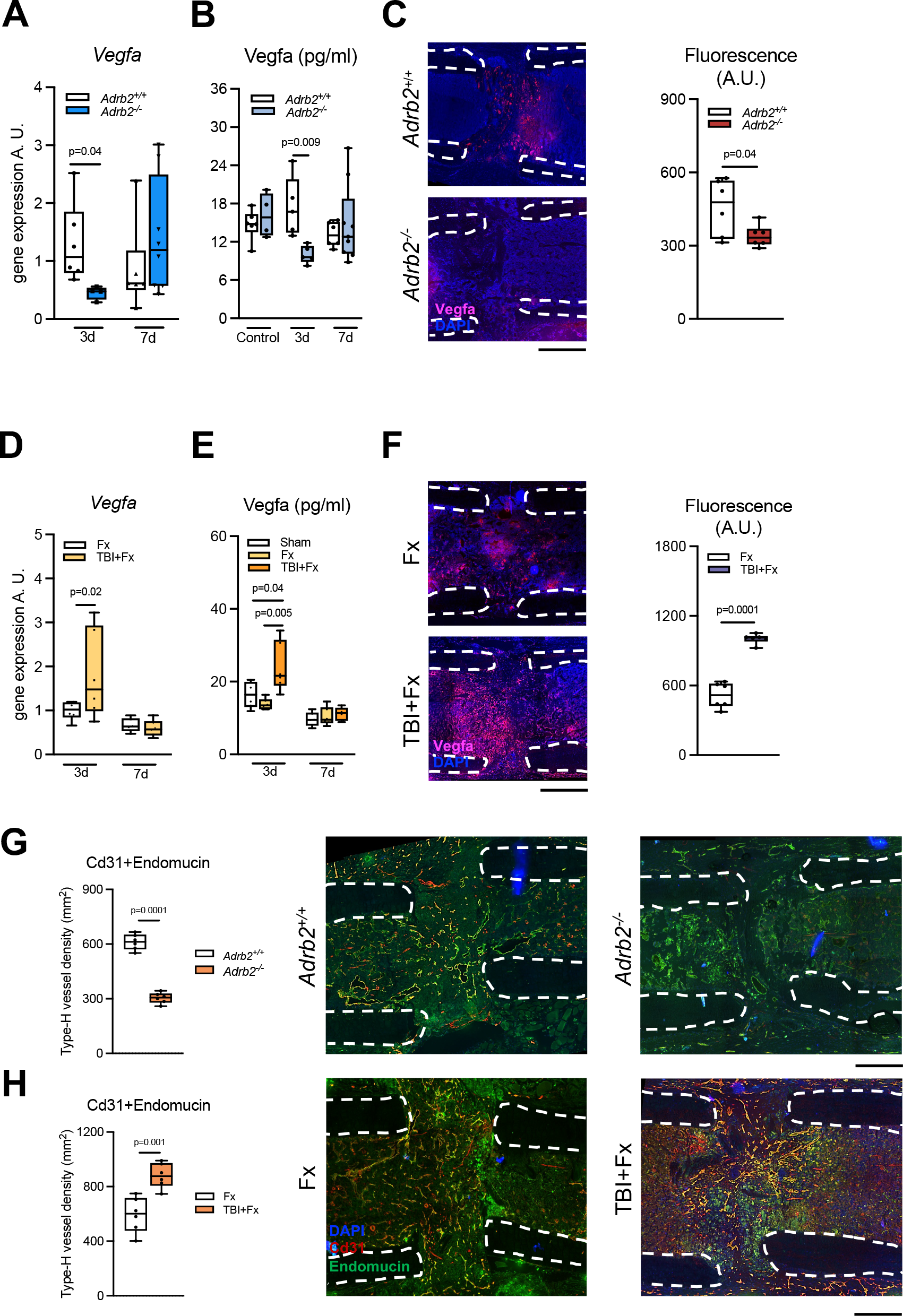
NE-Adrb2 signaling controls Vegfa levels and type-H vessel formation *in vivo*. (**A**) Expression of *Vegfa* in the callus 3 and 7 days after osteotomy in 30-week-old male mice of the indicated genotypes (FVB/129 genetic background). (**B**) Serum Vegfa levels in the same mice and untreated controls at the indicated time points. (**C**) Representative immunofluorescent stainings using a Vegfa-specific antibody in the fracture callus of 30-week-old male mice of the indicated genotypes 7 days after osteotomy, and quantification of signal intensity in the ROI. Scale bar=300 µm. (**D**) Expression of *Vegfa* in the callus 3 and 7 days after osteotomy in 12-week-old female WT mice (C57Bl/6J genetic background) with/without TBI. (**E**) Serum Vegfa levels in the same mice and sham-operated controls at the indicated time points. (**F**) Representative immunofluorescent stainings using a Vegfa-specific antibody in the fracture callus of 12-week-old female WT mice 7 days after osteotomy with/without TBI, and quantification of signal intensity in the ROI. Scale bar=300 µm. (**G**) Quantification of type-H vessel density and representative immunofluorescent co-stainings using Cd31- and Endomucin-specific antibodies in the fracture callus of 30-week-old male mice (FVB/129 genetic background) of the indicated genotypes 14 days after osteotomy. Scale bar=400 µm. (**H**) Quantification of type-H vessel density and representative immunofluorescent co-stainings in the fracture callus of 12-week-old female WT mice (C57Bl/6J genetic background) 14 days after osteotomy with/without TBI. Scale bar=400 µm. In (**C**), (**F**), (**G**), and (**H**), dotted white lines show the fracture ends. (**A**), (**B**), (**D**), and (**E**), two-way ANOVA followed by Tukey’s post hoc test. (**C**), (**F**), (**G**), and (**H**) two-tailed Student’s t-test. For all experiments, n=6 mice per group and time point. Numerical data are graphed in boxplots with median and 25^th^ and 75^th^ quantiles. Whiskers indicate upper and lower extremes, respectively.

Previously it was shown that Vegfa expression during bone regeneration is promoted by calcitonin-gene-related peptide (Cgrp), a neuropeptide also essential for an adequate fracture healing process (37, 38). Testing whether a similar mechanism might be present in periosteal cells, we found NE to increase *aCgrp* gene expression and Cgrp protein content in cell culture supernatants (**Figure 7A,B; Supplemental Figure 9A,B**). *In vivo*, *aCgrp* mRNA expression in the fracture callus was increased after TBI in the early healing phase, and associated with increased serum Cgrp levels 3 and 7 days following TBI (**Figure 7C,D**). Similarly, while serum Cgrp levels increased during fracture healing in WT mice, they remained unaltered in mice lacking Adrb2 (**Figure 7E**). These findings were confirmed locally in the fracture gap, where a reduced signal intensity of Cgrp immunofluorescence was observed in Adrb2-deficient mice 7 days after osteotomy, whereas the opposite was observed in fracture mice with concomitant TBI (**Figure 7F**). Mechanistically, Cgrp potentiated the stimulatory effect of NE on Vegfa expression in periosteal cells, and the Cgrp receptor antagonist, BIBN, inhibited the stimulatory effect of NE on Vegfa expression (**Figure 7G,H**). Likewise, NE failed to induce *Vegfa* and *Hif1a* expression in periosteal cells lacking *aCgrp* (**Figure 7I; Supplemental Figure 9C**). Similar observations were made in bone marrow-derived osteoblasts. Here, also less pronounced than in periosteal cells, NE induced *Vegfa* and *Hif1a* expression, which was blunted in osteoblasts derived from aCgrp-deficient mice (**Supplemental Figure 9D**). Finally, to test the functional relevance of these observations *in vivo*, aCgrp-deficient mice were subjected to TBI. Similar to Adrb2-deficiency, mice lacking aCgrp failed to respond to the beneficial impact of TBI on fracture healing as evidenced by radiological and histological outcome parameters (**Figure 7J,K**).

**Figure 7.**
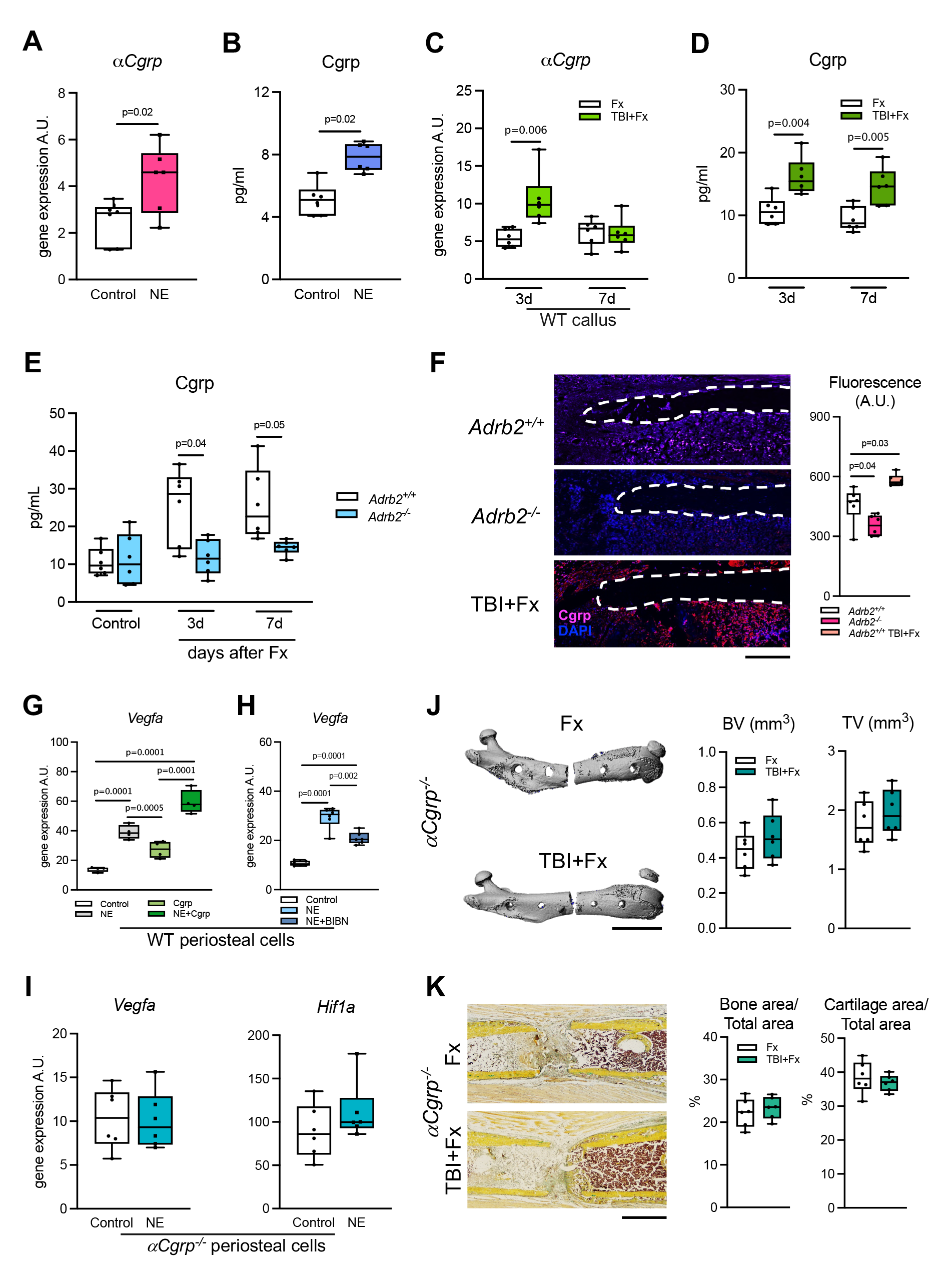
aCgrp-dependent induction of *Vegfa* by NE. (**A**) Expression of *aCgrp* by qRT-PCR in WT periosteal cells derived from femur after stimulation with NE for 6 hours and (**B**) Cgrp protein concentrations in the supernatant of the same cultures measured by ELISA. (**C**) Expression of *aCgrp* in the fracture callus of 12-week-old female WT mice (C57Bl/6J genetic background) with the indicated surgeries and (**D**) Cgrp concentrations in the serum of the same mice. (**E**) Cgrp concentrations in the serum of 30-week-old male mice (FVB/129 genetic background) of the indicated genotypes with/without a femoral osteotomy. (**F**) Representative immunofluorescent stainings using an Cgrp-specific antibody in the fracture callus of 30-week-old male mice of the indicated genotypes with a femoral osteotomy with/without TBI and quantification of signal intensity in the ROI 7 days after surgery. Dotted white lines show the fracture ends, scale bar=100 µm. (**G**) Expression of *Vegfa* in WT periosteal cells stimulated with NE, Cgrp, or both, for 6 hours. (**H**) Expression of *Vegfa* in WT periosteal cells stimulated with NE and/or the Cgrp receptor antagonist BIBN for 6 hours. (**I**) Expression of *Vegfa* and *Hif1a* in Cgrp-deficient periosteal cells stimulated with NE for 6 hours. (**J**) Representative µCT images of the fractured femora in 12-week-old female Cgrp-deficient mice (C57Bl/6 genetic background) with/without TBI 21 days following fracture and radiological quantification of callus bone volume (BV) and tissue volume (TV). Scale bar=5 mm. (**K**) Representative callus sections (Movat Pentachrome staining) and histomorphometric quantification of bone area/total area and cartilage area/total area of the fracture callus in the same mice. Scale bar=500 µm. (**A**), (**B**), (**I**), (**J**), and (**K**) two-tailed Student’s t-test. (**C**), (**D**) (**E**), (**F**), (**G**), (**H**) Two-way ANOVA followed by Tukey’s post hoc test. *In vitro*, n=6 independent cultures per group (derived from mice with C57Bl/6 genetic background). *In vivo*, n=6 mice per group and time point. Numerical data are graphed in boxplots with median and 25^th^ and 75^th^ quantiles. Whiskers indicate upper and lower extremes, respectively.

Together, these findings demonstrated that increased adrenergic signaling promotes fracture healing via an Adrb2-Vegfa axis, thus controlling the formation of type-H vessels in the callus and subsequently bone repair. To test this for therapeutic relevance, we next employed the clinically widely used, nonselective Adrb2 antagonist propranolol (blocking both beta1-and beta2 adrenoreceptors) (39) and the selective Adrb2 agonist formoterol, used as an inhaled agent in asthma or chronic obstructive pulmonary disease (COPD) (40), and assessed their impact on fracture healing in aged mice. Using µCT scanning, we found systemic propranolol treatment to impair the formation of new bone in the callus, whereas systemic formoterol application resulted in improved radiological outcome parameters (**Figure 8A**). The same observations were made on undecalcified cryosections of the fracture callus, where propranolol treatment resulted in a reduction of newly formed bone and formoterol increased mineralized callus volume (**Figure 8B**). Assessment of osseous callus bridging further demonstrated a high rate of fracture nonunion in mice treated with propranolol, and improved fracture union in mice receiving formoterol (**Figure 8C**). As an explanation, a decreased density of type-H vessels was detected in mice treated with propranolol, whereas the opposite effect was observed in the case of formoterol treatment (**Figure 8D; Supplemental Figure 10**). From a clinical perspective, we finally performed a retrospective cohort analysis of polytraumatized patients with long bone shaft fractures (humerus, femur, or tibia) and intravenous treatment with NE for cardiovascular support. Patients with TBI or spinal cord injury were excluded due to a confounding effect on fracture healing. As controls, we assessed otherwise healthy patients with a corresponding long bone fracture only (**Supplemental Table 1**). All patients received intramedullary nailing of the shaft fractures, and callus size was measured 6 months after surgery by two blinded investigators. Here, fracture patients that had received systemic NE treatment showed an increase in relative callus diameter, both in the anterior-posterior and sagittal plane (**Figure 8E**).

**Figure 8.**
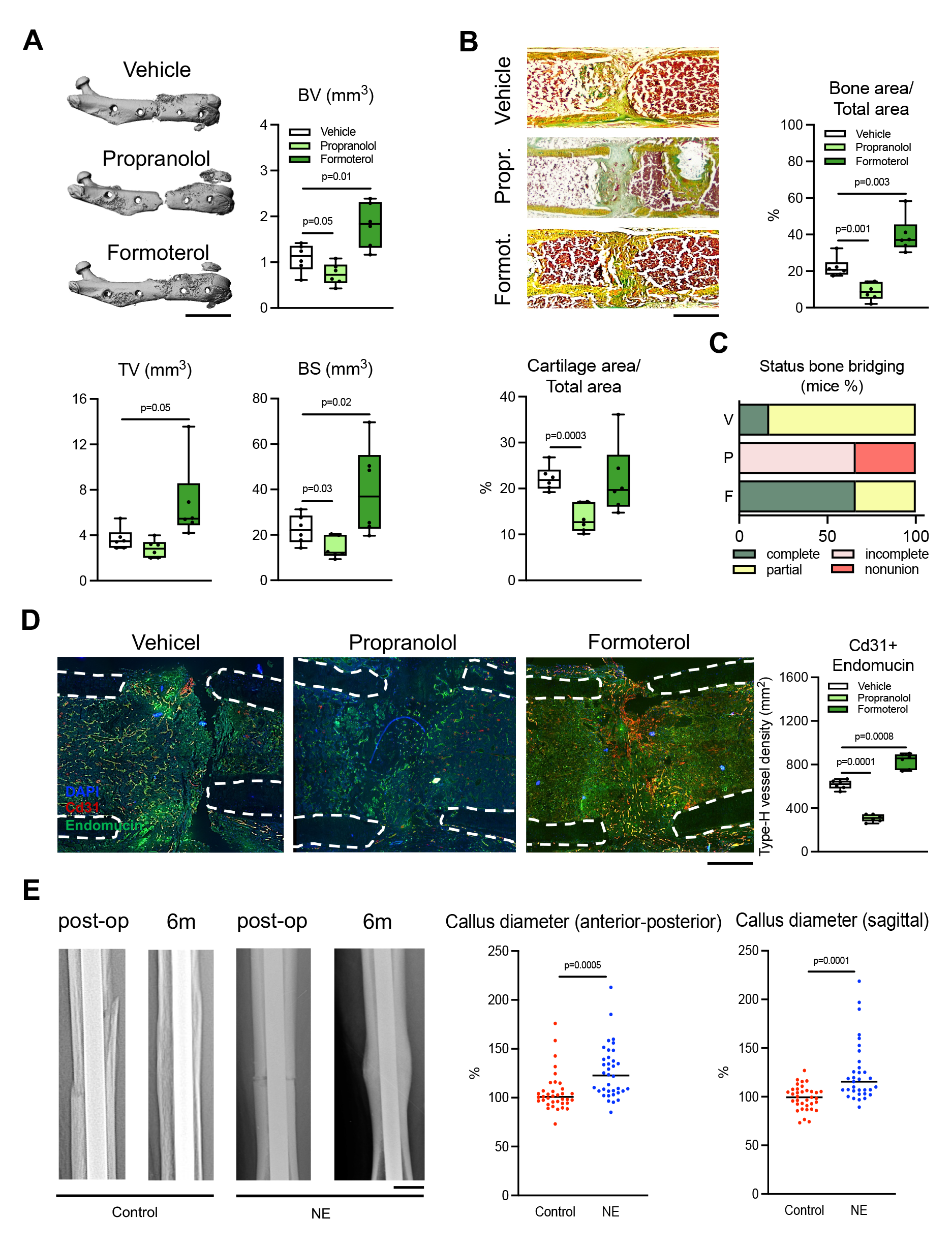
Modulation of fracture healing by Adrb2 agonism and antagonism, respectively. (**A**) Representative µCT images of the fractured femora in 30-week-old male WT mice (FVB/129 genetic background) treated daily intraperitoneally with vehicle, propranolol, or formoterol during the entire study period (21 days) and quantification of callus bone volume (BV), tissue volume (TV), and bone surface (BS) in the callus. Scale bar=5 mm. (**B**) Representative callus sections in the same mice and quantification of bone area/total area and cartilage area/total area. Scale bar=500 µm. (**C**) Semiquantitative evaluation of osseous callus bridging in the same mice. (**D**) Representative immunofluorescent co-stainings using Cd31- and Endomucin-specific antibodies in the fracture callus of the same mice and quantification of type-H vessel density. Dotted white lines show the fracture ends, scale bar=200 µm. For (**A**)-(**D**), two-way ANOVA followed by Tukey’s post hoc test. n=6 mice per group and time point. Numerical data is graphed in boxplots with median and 25^th^ and 75^th^ quantiles. Whiskers indicate upper and lower extremes, respectively. (**E**) Representative radiographs of tibial fractures from patients with/without systemic NE treatment immediately post-operatively and 6 months (6m) after intramedullary nailing (scale bar=15 mm), and quantification of relative callus diameter in anterior-posterior and sagittal plane of the healed fractures. n=37 patients per group, two-tailed Student’s t-test.

## Discussion

In the search for novel approaches to pharmacologically improve bone healing and prevent fracture nonunion, the current study investigated the unique clinical phenomenon that TBI improves fracture healing while inducing bone loss in the uninjured skeleton (14). Our data show that increased NE-Adrb2 signaling mediates TBI-induced skeletal manifestations in terms of both bone remodeling and regeneration. Furthermore, the study demonstrates that Adrb2 plays an age-dependent and pharmacologically targetable role in fracture healing by promoting callus vascularization under conditions of increased sympathetic tone.

The clinical observation that TBI accelerates bone healing has attracted the interest of many researchers as it may hold the key to develop novel treatment approaches to improve fracture healing (14). This is particularly important because, unlike osteoporosis where antiresorptive and osteoanabolic agents are readily available, there are no pharmacologic options that reliably improve bone healing and prevent nonunion (4, 7). Intermittent injections of teriparatide, a fragment of human parathyroid hormone that promotes bone formation, have shown promising results in animal models, but compelling clinical studies supporting these observations are lacking (41). In addition, although topical application of bone morphogenetic proteins can potently stimulate osteogenesis and thus bone repair, their use is limited by serious adverse effects, including inflammatory tissue swelling with compression of nerve structures, ectopic bone formation, excessive osteolysis, and wound healing complications (42).

Because callus is a very heterogeneous tissue consisting of a dynamically changing composition of many different cell types during overlapping healing stages, our previous attempts to decipher the molecular signature of bone regeneration after TBI have been largely inconclusive. Thus, we took advantage of the fact that TBI also affects bone remodeling in unfractured bone, which is much more homogeneous in tissue composition compared to callus. Unbiased RNA-seq not only showed reduced expression of osteoblast markers after TBI but also revealed suppressed expression of *Adrb2* which was confirmed in independent experiments via qRT-PCR. Since Adrb2 is a classic example of a G-protein coupled cell surface receptor that is downregulated at the mRNA level upon excessive stimulation with endogenous ligands (24), including NE, these findings suggested a hyperadrenergic state in the TBI model, similar to that observed in patients. This was confirmed by increased serum normetanephrine levels and NE content in the femora of TBI mice. In this regard, previous seminal work identified Adrb2, one of the major receptors for NE, as an essential regulator of both osteoblast and osteoclast function (2, 31, 43). Stimulation of Adrb2 by NE inhibits osteoblast proliferation and bone formation (44). In addition, activation of Adrb2 in osteoblasts results in increased bone resorption through induction of RANKL, the master paracrine regulator of osteoclastogenesis (45). In our study, these previously described effects of NE-Adrb2 signaling were clearly mirrored in the unfractured skeleton of TBI mice, which exhibited reduced bone volume due to decreased bone formation indices and increased osteoclast parameters in the face of a hyperadrenergic state. Mechanistically, these skeletal changes were abolished in Adrb2-deficient mice subjected to TBI, providing genetic evidence that, in the unfractured skeleton, TBI leads to the deterioration of bone remodeling via Adrb2.

In terms of fracture healing, the beneficial effect of TBI on bone regeneration was also completely absent in mice lacking Adrb2, demonstrating an essential function of Adrb2 in promoting bone repair in this condition. This was indeed surprising, as previous studies have reported inconsistent results regarding the role of beta-adrenergic signaling in bone repair, including no effect or even impaired healing. For example, Minkowitz et al. reported that propranolol improved the fracture union of bone defects filled with demineralized bone matrix powder in young rats (46). However, this finding was not confirmed in an independent study, in which propranolol did not improve or inhibit the healing and mechanical strength of osteotomies in rats of unreported age (47). Another study showed that local, but not systemic, inhibition of beta-adrenergic signaling stimulated the healing of bone defects in rats of unspecified age (48). Finally, a more recent study suggested that a single preoperative injection of propranolol improved bone healing in a mouse model of chronic psychosocial stress by modulating immune cell function (49).

Therefore, to better understand the role of Adrb2 in bone repair, we examined fracture healing in young and otherwise healthy Adrb2-deficient mice without TBI. Despite a trend toward increased bone volume in the early stages of healing observed only histologically, but not radiographically, normal callus mineralization and fracture union were observed in young Adrb2-deficient mice 21 days after osteotomy. In sharp contrast, aged Adrb2-deficient mice exhibited severely impaired bone healing and a high rate of fracture nonunion with only limited callus remodeling by osteoblasts and osteoclasts. As an explanation for this age-dependent role of Adrb2 in bone regeneration, we detected increased NE content in the bones of aged mice, confirming previous studies demonstrating low local NE reuptake in cortical bone and consequently enhanced NE signaling (33). These findings are also consistent with an overall increased sympathetic tone in the aged organism, which is well-established not only in animal models but also in humans (32). Thus, while only of minor significance to bone repair in young and healthy mice, potentially explaining inconsistent results of aforementioned previous studies, Adrb2 essentially controls fracture healing under conditions of increased sympathetic tone, including TBI and aging.

Since the effects of Adrb2 on bone healing in TBI or aged mice could not be explained by the established inhibitory function of Adrb2 on osteoblastic bone formation, we monitored the effects of NE stimulation on periosteal cells *in vitro*, which express high levels of Adrb2 in cortical bone *in vivo*. While NE, unlike in osteoblasts, did not affect their osteogenic capacity, it caused a potent, Adrb2-dependent induction of the angiogenic factors *Vegfa* and *Hif1a*. Both peptides regulate the formation of type-H blood vessels, which are essential for new bone formation during fracture healing (35, 36). Consequently, while the callus of TBI mice showed an increased density of type-H vessels, aged Adrb2-deficient mice exhibited the opposite vascular phenotype, suggesting that enhanced NE-Adrb2 signaling is a critical regulator of callus vascularization.

The importance of callus vascularization in bone regeneration is best illustrated by several clinical studies that clearly demonstrate that vascular disease is a major risk factor for fracture nonunion (35, 50). Similarly, damage to the periosteum, whose integrity is a prerequisite for sufficient callus vascularization, impairs bone healing (51). In this regard, the neuropeptide Cgrp is known to be highly expressed in the periosteum and essential for adequate fracture healing (38). As a previous study described a Cgrp-dependent induction of *Vegfa* expression during bone healing, we also tested the functional role of Cgrp in the stimulatory effect of NE on *Vegfa* expression (37). Like *Vegfa*, *aCgrp* was induced by NE in periosteal cells and to a lesser extent also in osteoblasts, whereas aCgrp-deficient mice did not respond to the beneficial effects of TBI. Similarly, aCgrp-deficient cells failed to respond to stimulation of *Vegfa* expression by NE, and co-treatment of NE and Cgrp potentiated the stimulatory effect on *Vegfa* induction. Since *Vegfa* induction by NE was blunted by the Cgrp receptor antagonist BIBN, these observations suggest an autocrine mechanism by which Cgrp modulates the stimulatory effect of NE on *Vegfa* expression. Although our study focused primarily on periosteal cells, we also found that NE-Adrb2 signaling increased *Vegfa* expression in osteoblasts. Thus, it is likely that increased NE signaling induces the production of angiogenic factors not only in periosteal cells but also in other cell types during bone repair, as Adrb2 has been repeatedly shown to be a positive regulator of angiogenesis in many organ systems (52). For example, it has been demonstrated that NE dose-and time-dependently induces *Vegfa* transcription in brown adipocytes, which is reversible by the beta-blocker propranolol (53, 54). Similarly, adrenergic signaling through Adrb2 has been shown to promote Vegfa-dependent tumor vascularization in prostate, skin, and breast cancer cells (55-57). Taken together, these observations demonstrate that NE is capable of inducing Vegfa in a variety of different cell types and, in the case of bone healing, results in an overall promotion of type-H vessel formation required for osteogenesis.

Since key observations in this study were made in both female and male mice with C57Bl/6J or FVB/129 genetic backgrounds, it is likely that the identified mechanism of NE-Adrb2 action in bone regeneration may also be relevant in other species, including humans. From a translational perspective, the above findings in mice are supported by our clinical evaluation of fracture patients treated with intravenous administration of NE. These patients received systemic NE for cardiovascular support after polytrauma with life-threatening injuries, which typically worsens fracture healing outcomes (58). Therefore, the association of systemic NE treatment with increased bone callus formation six months after injury is indeed striking. Due to the retrospective nature of this clinical study, we additionally treated aged WT mice with the selective Adrb2 agonist formoterol after inducing osteotomy. Systemic application of this drug, which is widely used clinically as an inhaled agent for asthma and COPD, promoted type-H vessel formation and improved fracture healing, demonstrating that pharmacological Adrb2 agonism is a suitable approach to stimulate bone regeneration, at least in mice.

While these data identify the cell surface receptor Adrb2 as a novel target for promoting bone healing, they also suggest that its blockade may result in impaired fracture healing. Indeed, we additionally found that the nonselective beta-blocker propranolol causes delayed fracture healing in aged mice. Our findings are essentially confirmed by the independent study by Steffenson et al., who observed that beta-blocker exposure was associated with nonunion in 253,266 fracture cases (medRxiv/2023/292608, Lillia Steffenson and Lucas Marchand, University of Utah, USA, personal communication). This is of high clinical relevance, as > 22 % of people aged 60 years or older are reported to be taking beta-blockers (59). Based on the previously described beneficial effects of beta-adrenergic antagonism in the unfractured skeleton, an NIH-funded randomized controlled trial is currently investigating the effect of beta-blockers in the prevention of osteoporosis (60). In this ongoing trial, but also in everyday clinical practice, our observations imply that a temporary discontinuation of beta-blockers in elderly patients after a fracture event should be considered, if of course justified from a cardiovascular point of view.

Taken together, our results are unexpected and propose a novel function of Adrb2 signaling in the injured skeleton. They show that under conditions of increased sympathetic output, such as in TBI or, most commonly, as a physiological adaptation in the aging organism (32), Adrb2 signaling is an essential stimulus for bone healing. They also help to explain why, contrary to popular belief, fracture healing is not impaired in healthy elderly individuals, who exhibit reduced rates of fracture nonunion compared to young patients with lower sympathetic tone (61-63). As the incidence of age-related fractures is steadily increasing (6) and a significant proportion of those affected are exposed to beta-blockers (59), these findings are of high translational relevance, and clinical studies are warranted to test their significance in clinical settings.

## Methods

### Animals

Wild-type (WT) mice with FVB/129 or C57Bl/6J background were used as indicated in the figure legends. Adrb2-deficient mice (*Adrb2tm1Bkk/J*; Jackson Lab #031496) exhibited FVB/129 genetic background, and aCgrp-deficient mice (*Calcatm1Eme;* MGI #21485389) exhibited C57Bl/6J genetic background (both lines backcrossed at least 7 times) (64). Animals were maintained under standard conditions (12 hour light-dark circadian rhythm, 22 °C) in an SPF facility and housed in stable groups. Water and a standard diet were provided *ad libitum*. For all animal experiments, the directly compared groups like sham, isolated fracture, or fracture in combination with TBI were performed in parallel and treated equally throughout the experiment. The *a priori* group size calculation for fracture healing evaluation was n=6 mice - the exact number of analyzed animals is given in the figure legends. Littermates were randomized to the different experimental groups and all biological samples were coded to facilitate blinded evaluation. Both male and female mice were used for the experiments as indicated in the figure legends.

### Surgery

Mice received a femoral osteotomy stabilized with an external fixator as previously described. Briefly, mice were anaesthetized, and the right femur was exposed. An external fixator (RISystem, Davos, Switzerland) was mounted on the femur, using four pins and a 0.45 mm hand drill. The fracture gap was created using a 0.7 mm diameter Gigli wire saw. The wound was closed with a simple interrupted suture. TBI to the left parietotemporal cortex was induced as previously described (22). Following the craniotomy, a controlled cortical impact (bolt 3 mm flat, 45 ° angle, impact velocity of 3.5 m/s, 0.25 mm penetration depth, contact duration of 0.15 s) was induced on the intact dura mater. Subsequently, the skull piece was repositioned, fixed with dental cementum, and the skin incision was closed with Ethilon 6.0 suture. For sham surgery, the same protocol was followed, except for the controlled cortical impact onto the dura mater. For the combination of both injuries, TBI was induced prior to the fracture. Clindamycin (150 mg·kg-1, Hikma Pharma GmbH) and buprenorphine (0.1 mg·kg-1, Richter Pharma AG) were administered preoperatively. For optimal pain relief, metamizole (1 mg·m-1, Novaminsulfon-ratiopharm®) or tramal (0.3 mg·ml-1, Grünenthal) was administered in the drinking water for 3 days postoperatively.

### Pharmacological treatment

From the time point of surgery until sacrificing, mice received two intraperitoneal injections daily of either vehicle (0.9% saline), formoterol fumarate dihydrate (10 mg·kg^-1^, Merck) or propranolol hydrochloride (20 mg·kg^-1^ for the first five days and 10 mg·kg^-1^ from day six until euthanasia, Merck), diluted in 0.9% saline, with a total volume of 100 μl intraperitoneally. Mice were sacrificed on postoperative day 14 or 21 as indicated.

### Micro-computed tomography

Fractured femora were fixed in 4 % paraformaldehyde at 4 °C for 24 h. After removing the fixators the bones were washed three times and stored in PBS during μCT scanning. Both fractured and unfractured femora were scanned using μCT (VivaCT 80, SCANCO Medical AG, Brüttisellen, Switzerland) with a voxel resolution of 15.6 μm, 400 ms integration time, 70 kVp and 113 μA. The callus evaluation was performed in a volume of interest (VOI) of 1 mm x 2 mm x 2 mm around the fracture gap. Data are presented according to the American Society for Bone and Mineral Research (ASBMR) guidelines for μCT analysis (66). For evaluation of unfractured femora, an 1-2 mm mid-shaft and 1-2 mm distal segment were analyzed. Cortical and trabecular properties were derived using an automated image analysis algorithm provided by the manufacturer.

### Biomechanics

Following µCT evaluation, all bones harvested on day 28 were challenged with a destructive three-point bending test as previously described (67). In brief, femora were placed on two support bars of the device (ZwickRoell) and a bending load with a maximum of 20 N was applied medially onto the callus site. Derived parameters were calculated automatically by the device software.

### Cryo-Embedding

After µCT scans, the bones were incubated in ascending sugar gradient (10 %, 20 %, and 30 % each for 24 h at 4 °C) and embedded in SCEM medium (Section Lab Co Ltd.), frozen in hexane (Carl Roth GmbH&CoKG) and stored at -80 °C until further processing. Using a cryostat (Leica CM3050S, Leica Microsystems), longitudinal sections of 5-7 µm were derived and mounted onto microscope slides using cryofilm (Cryofilm type II C, Section Lab Co Ltd.).

### Immunofluorescence staining

For Cgrp-specific staining, frozen femur sections were blocked with 3% BSA and 5% goat serum in PBS, incubated with Cgrp antibody (rabbit polyclonal, abcam #ab47027, 1:300 in DAKO antibody diluent (Agilent, #S0809)), overnight at 4 °C, washed three times in PBS and incubated with the secondary antibody (donkey anti-rabbit Cy3, Dianova, #711-165-152, 1:400 in 3% BSA and 5% donkey serum in PBS) for 1 hour. After washing with PBS and distilled water, slides were mounted with Fluoromount and DAPI (Southern Biotech, # 0100-20) and stored at 4 °C. For Adrb2-specific staining sections were fixed for 5 min at 4 °C with ice-cold acetone, permeabilized with Proteinase K (0,24 U/ml) and incubated with Adrb2 antibody (rabbit polyclonal, Bioss Antibodies # bs-0947R, 1:250) overnight at 4 °C. For Vegfa and vessel staining, sections were blocked in 3% BSA and 5% donkey serum. The primary antibodies (anti-Cd31, goat polyclonal, R&D Systems #AF3628, 1:100; anti-VEGFA, rabbit monoclonal, abcam #ab52917, 1:200; anti-Endomucin, rat, Santa Cruz #sc-65495, 1:100 in AK Diluent) and secondary antibodies (anti-goat 546, Thermo Fisher, #A-11058, 1:400; Alexa Fluor 488 anti-rabbit, Thermo Fisher, #A-21206, 1:400; Alexa Fluor 546 anti-rabbit, Thermo Fisher, #A-11056, 1:400; anti-rat 488, Thermo Fisher, #A-21208, 1:400) were used. For all immunofluorescent stainings, images at the indicated magnifications were acquired using the Leica DM RB microscope (Leica Microsystems) and Axiovision Rel. 4.8 software (Zeiss) as well as LEICA SP5 confocal microscope equipped with a Mai Tai HP multiphoton laser (Spectra Physics). Signal intensities in the ROI of the healing fracture (1 mm x 2 mm square placed centrally over the fracture gap) were quantified by two blinded investigators using ImageJ Analysis System (NIH).

### Histological stainings

Movat’s Pentachrome staining was performed as described previously (68). For this, sections were stained in alcian blue (8GS, Chroma), Weigert’s haematoxylin (Merck), Brilliant Crocein/Acid Fuchsin (Brilliant Crocein R, Chroma, and Acid Fuchsin, Merck), 5% Phosphotungstic acid PTA (Chroma), and Saffron du Gâtinais (Chroma). The stained sections were mounted in Vitro-Clud (Langenbrinck). For histomorphometry, mosaic images were taken using the Axioscop 40 (Zeiss) and Axiovsion Rel.4.8 software. Static and cellular histomorphometry were performed in the region of interest (ROI; 1 mm x 2 mm square placed centrally over the fracture gap) of one section per animal and time point. The distribution of cartilage and mineralization within the callus area was calculated using ImageJ Analysis System (NIH). Callus bridging of these specimens was assessed by two blinded investigators and categorized using the following scoring system: a=complete bridging (all four cortices bridged by callus), b=partial bridging (two or three cortices bridged by callus), c=incomplete bridging (callus present but no bridging visible), and d=nonunion (rounded cortices, minimal presence of callus). For TRAP activity staining, sections were fixed with 4% PFA and titrated to pH 5 using a buffer containing sodium acetate (Merck) and sodium tartrate-dehydrate (Merck). After washing with distilled water, sections were stained in the presence of Naphtol AS-MIX-Phosphate, Fast Red Violet LB Salt, N,N-Dimethylromaid and Triton X (Sigma Aldrich). A counterstain was carried out with Mayer’s Hemalaun solution (Merck). Osteoclasts were identified as TRAP-positive cells with ≥3 nuclei and adherent to the bone surface. Osteoblasts were identified as mononuclear cells adherent to the bone surface with typical cuboidal morphology. For the histomorphometry of the unfractured spine and femur the whole skeleton was fixed in 4 % formaldehyde for 48 h, vertebra bodies L1-L4 and the unfractured femur were dissected, dehydrated, and embedded undecalcified in methylmetacrylate as previously described (69). 4 μm sections of the spine and the femur, respectively, were cut using a Microtec rotation microtome. Von Kossa/van Gieson, TRAP-activity, and Toluidine blue staining were performed as described before (69). For the evaluation BioQuant and the OsteoMeasure histomorphometry system (Osteometrics Inc.) were used.

### Calcein labeling of bone formation

Prior to skeletal analysis (14 and 21 days postoperatively), mice were injected twice with calcein (30 mg kg^−1^ body weight, Sigma C-0875) 9 and 2 days before sacrifice. Calcein-positive area per total callus area was calculated as a surrogate marker for newly formed bone in the callus using ImageJ Analysis System (NIH).

### RNA extraction and qRT-PCR

The callus, including both fracture ends, was extracted between the two central pins. Midshaft femoral sections were dissected and served as unfractured control samples. Following snap freezing in liquid nitrogen, further processing was carried out using a standardized purification protocol employing homogenization in TRIzol (Sigma Aldrich, Merck) with an Ultra Turrax (IKA Labortechnik). For isolation of RNA from periosteal cells and osteoblasts, cells were washed and lysed in TRIzol (Sigma Aldrich, Merck). Total RNA of all samples was then isolated using columns of a NucleoSpin RNA kit (Macherey-Nagel). The concentration and quality of the derived RNA were monitored by using NanoDrop 2000 system (NanoDrop Technology). Reverse transcription into cDNA was performed using a cDNA synthesis kit (ProtoScript First Strand cDNA Synthesis Kit, New England BioLabs, Ipswich) according to the manufacturer’s instructions. Quantitative real-time PCR (qRT-PCR) was conducted using TaqMan Assay-on-Demand primer sets (Applied Biosystems by Thermo Fisher Scientific) or Power SYBR Green PCR Master Mix (Merck). For all groups, values are displayed as an expression over the housekeeping gene Glyceraldehyde-3-phosphate dehydrogenase *(Gapdh)* as a reference. The following primers were used for TaqMan: *Acp5* (Mm00475698_m1), *Alpl* (Mm00475834_m1), *Bglap* (Mm03413826_mH), *Calcr* (Mm00432271_m1), *Clcn7* (Mm00442400_m1), *Col1a1* (Mm00801666_g1), *Ctsk* (Mm00484039_m1), *Dmp1* (Mm01208363_m1), *Gapdh* (Mm99999915_g1), *Hif1a* (Mm00468869_m1)*, Ibsp* (Mm00492555_m1), *Phex* (Mm00448119_m1), *Postn* (Mm00450111_m1), *Rankl = Tnfsf11* (Mm00441906_m1), *Runx2* (Mm00501580_m1), *Sost* (Mm00470479_m1)*, Sp7* = *Osx* (Mm00504574_m1), *Vegfa* (Mm00437306_m1), *Gapdh* (Mm99999915_g1). Primers for SYBR Green were designed as follows: *Adrb2* (FOR-CCATGACTTCATTGCACCCG, REV-CACTCATCGGTCACGACACA), *aCgrp (*FOR-CAGGCCTGAACAGATAACAGC, REV-TGTGTCTTTCATCAGCCTTTCTT), *Gapdh (*FOR-TGCACCAACTGCTTAG, REV-GGATGCAGGGATGATGTTC), *Slc6a2 (*FOR*-*TGTGTTTGTACGCCCAGGTG, REV*-*ACCTCCTAGCACCTTCGTGA). Gene expression was calculated by the ΔΔCT method as virtual copy number per housekeeper gene (arbitrary units, A.U.) or as fold expression as indicated.

### RNA-Sequencing

n=3 (sham) and n=4 (TBI) unfractured femur samples were employed. For whole-exome sequencing, libraries were prepared using the TruSeq Stranded mRNA Kit (Illumina). Sequencing was performed by NovaSeq 6000 (Illumina) with 2 x 150 bp and 60 Mio reads per sample. RNA-seq reads were mapped to the mouse genome (GRCm38, version M12 (Ensembl 87)) with STAR version-2.7.9a (70) using the following parameters -quantMode GeneCounts –outSAMunmapped Within -outFilterType BySJout -outFilterMultimapNmax 20 - alignSJoverhangMin 8 -alignSJDBoverhangMin 1 -outFilterMismatchNmax 999 -outFilterMismatchNoverLmax 0.04 -alignIntronMin 20 -alignIntronMax 1000000 -alignMatesGapMax 1000000. Reads were assigned to genes using FeatureCounts SUBREAD version-v2.0.1 (71) with the following parameters: -T 2 -t exon -g gene_id -s 2 -p. For the differential expression analysis, we used DESeq2 version-1.32.0 (72) with default parameters. We filtered genes that had less than 5 counts in at least 3 samples. The gene set enrichment analysis was carried out using the CERNO algorithm from the R tmod package version-0.50.06 (73).

### ELISA

Measurements were conducted according to the protocols provided by the manufactures (Cgrp: Cloude Clone Corp., CEA876Mu; Vegfa: Quantikine, R&D Systems Biotechne Brand, MMV00; Normetanephrine: Demeditec Diagnostics, DEE8200).

### Norepinephrine content in bone

Measurement of NE in bone was performed similarly as described previously (33). In brief, femora were cleaned of connective tissues and the epiphyses including the articular cartilage were removed. Bone marrow was removed from cortical bone via centrifugation at 3000*g* for 2 min, and bones were then snap-frozen in liquid nitrogen. Frozen bones were crushed in a mortar and pestle, and resuspended in 100 μl of 0.1% reduced l-glutathione (Sigma-Aldrich) in 0.1 M Tris pH 8.0 to protect from oxidation. NE levels were measured by ELISA (Immusmol, BA-E-5200R) according to the manufacturer’s protocol. An aliquot of tissue suspended in extraction buffer was used to quantify protein concentration for normalization of NE measurements.

### Cell culture

Female and male WT and mutant mice at the age of 12-18 weeks with C57Bl/6J or FVB/129 genetic background were used as indicated. Bone marrow-derived osteoblasts were obtained by flushing out the bone marrow, filtering it through a 70 µm cell strainer, and centrifugation at 1000 RPM for 10 minutes. For periosteal cell culture, epiphyses of the femora were removed, and bones were flushed out several times to remove bone marrow. In the case of calvariae, bones were dissected free of connective tissue. Digestion of periosteal tissue from femora and calvariae was then performed by incubation with 0.1 % dispase (Roche Diagnostics GmbH) and 0.25 % collagenase II (Gibco Life Technologies Corporation) at 37 °C for 3 h. Periosteal cells were separated using a 70 µm cell strainer and centrifugation at 1000 RPM for 10 minutes. Subsequently, cells were plated onto 12-well plates in a concentration of 1 × 10^4^ cells per well. α-MEM (Merck KGaA) supplemented with 10 % FCS was used for cultivation and changed every 2-3 days. Osteoblast differentiation was induced through stimulation with 25 μg·mL−1 ascorbic acid and 5 mmol·L-1 β-glycerophosphate when cells had reached 70% confluence for 10 days. For alizarin red staining, cells were incubated with 40 mM alizarin red staining solution (pH 4.2) for 10 min at room temperature after fixation in 90% ethanol. To quantify alizarin red incorporation, cells were washed with PBS and fixed in 90% ethanol for 1 h. Following additional washing, the cell-bound alizarin red was dissolved in 10% acetic acid. After incubation for 30 min at room temperature and 10 min at 85 °C, the supernatant of a subsequent centrifugation step was neutralized with 10% ammonium hydroxide solution, and the absorbance was measured at 405 nm. For short-and long-term stimulations of periosteal cells and osteoblasts, the following conditions were employed: NE (Arterenol, Chelapharm, #03870227, 10^-5^ M), BIBN 4096 (Tocris #4561, 10^-3^ M), recombinant mouse Cgrp (Bachem #H-2265, 10^-7^M).

### Clinical study

Medical records of all patients admitted to the largest German level I trauma center (Charité – Universitätsmedizin Berlin, Campus Virchow Klinikum, Center for Musculoskeletal Surgery, Berlin, Germany) and diagnosed with a traumatic shaft fracture of the humerus, femur, or tibia (A1-A3 and B1-B3 according to the AO surgical reference) between 2008 and 2022 were identified by patient management software (SAP Business Client 6.5, SAP Walldorf). Inclusion criteria were: traumatic midshaft fractures of the humerus, femur, or tibia that were surgically stabilized with an intramedullary nail and recorded radiographic follow-up for six months. Exclusion criteria were: pathologic fractures, no six-month radiographic follow-up, no recorded ISS, traumatic brain injury, or spinal cord injury. A total of 850 patients were identified. 57 patients were excluded because of concomitant traumatic brain injury or spinal cord injury. 605 patients were excluded because they were lost to follow-up or had pathologic fractures. 188 patients were finally identified, of whom 37 had received NE therapy during their ICU stay and 151 had not. Propensity score matching for age, Injury Severity Score (ISS), sex, and American Society of Anesthesiologists (ASA) score was performed using "R", the MatchIt package, and the RStudio© software (RStudio, Inc.) to create balanced and homogeneous groups. This resulted in a total of 74 patients, 37 patients with and 37 patients without systemic NE therapy (n=3 humerus fractures, n=16 femoral fractures, n=18 tibial fractures per group). Fracture healing was analyzed radiologically by measuring the maximum diameter of callus formation on anterior-posterior and axial radiographs six months postoperatively using radiographic calibration markers. The measurements were then divided by the maximum diameter of the fracture site measured on radiographs one week after surgery to calculate the resulting relative increase in callus growth.

### Statistical analysis

Sample sizes in this study were calculated based on our laboratory’s prior work according to the method provided in “http://www.lasec.cuhk.edu.hk/sample-size-calculation.html” with a statistical power of 80% and a level of significance of 0.05. For all *in vivo* experiments, n=6 mice per group and time point were used. For *in vitro* studies, n=6 independent cell cultures were employed. All groups were assigned randomly, and researchers were blinded during sample processing and analyses of outcome measurements. Unpaired two-tailed Student’s t-test was used to determine simple comparisons. For multiple-group comparisons, data were analyzed by one-way or two-way ANOVA test as indicated, followed by Tukey’s post-hoc tests. Numerical data are graphed in boxplots with median and 25^th^ and 75^th^ quantiles, whiskers indicate upper and lower extremes, respectively. The significance level was set at p < 0.05 (GraphPad Prism 9.1.1, La Jolla).

### Study approval

All experimental procedures were approved by the local legal representative animal rights protection authorities (Landesamt für Gesundheit und Soziales Berlin, G0009/12; G0007/17; G0147/18; G0112/22 and Behörde für Justiz und Verbraucherschutz Hamburg, N085/20; N090/21) and performed according to the principles established by the national institute of health guide for care and use of laboratory animals as well as the Welfare Act (Federal Law Gazette I, p.1094). The anonymized retrospective clinical study was approved by the local ethics committee (EA4/038/17).

### Data availability

The individual data points are displayed in the figures and underlying data is also available as an Excel file linked to the manuscript. Furthermore, exome sequencing raw data is publicly available and further material will be provided upon reasonable request.

## Author contribution

J.S., C.G., M.F., S.J., M.R., C.E., J.A., J.W., A.R., W.X. and A.D. conducted specific experiments and analyzed specific data. F.G., L.R. and Q.K. analyzed patient data. A.I., M.N.T., and D.B. analyzed RNA-seq data. G.D., T.S., K-H.F., S.T. critically revised the manuscript. A.B. and J.K. wrote the manuscript. J.K. and S.T. designed the research studies. D.J., P.R.K., E.O. and P.K. share the first author position. D.J. was responsible for all animal experiments combining TBI and fracture, comprising WT, aCgrp-deficient and Adrb2-deficient mice. She collected and processed the samples and did the majority of the data analysis. P.R.K. was responsible for all animal experiments of isolated fracture healing in Adrb2-deficient mice and respective WT cohorts. He collected and processed those samples and did the majority of data analysis, performed staining of the vasculature and *in vitro* analysis. Furthermore, he identified the age dependency. E.O. took care of experiments in WT and aCgrp-deficient mice, was essential for sample processing and evaluation of gene expression and histology. P.K. performed the fracture surgeries in the combined injury models, analyzed gene expression and gave valuable scientific input to the research direction. All authors reviewed and approved the final version of the manuscript.

## Acknowledgements

We would like to thank Olga Winter and Andrea Thieke for their assistance with the undecalcified histology of unfractured bone samples. This work was funded by grants from the German Research Foundation to J.K. (KE 2179/2-1, KE 2179/2-3) and S.T. (TS 303/2-1, TS 303/2-3, Project-ID 427826188 - SFB 1444). D.J. received funding from the Einstein Center Regenerative Therapies of Charité-Universitätsmedizin Berlin. P.R.K. is funded by a scholarship from the Werner-Otto Foundation, Hamburg, Germany.

