## Supplemental Information for "Increased beta2-adrenergic signaling is a targetable stimulus essential for bone healing by promoting callus neovascularization"

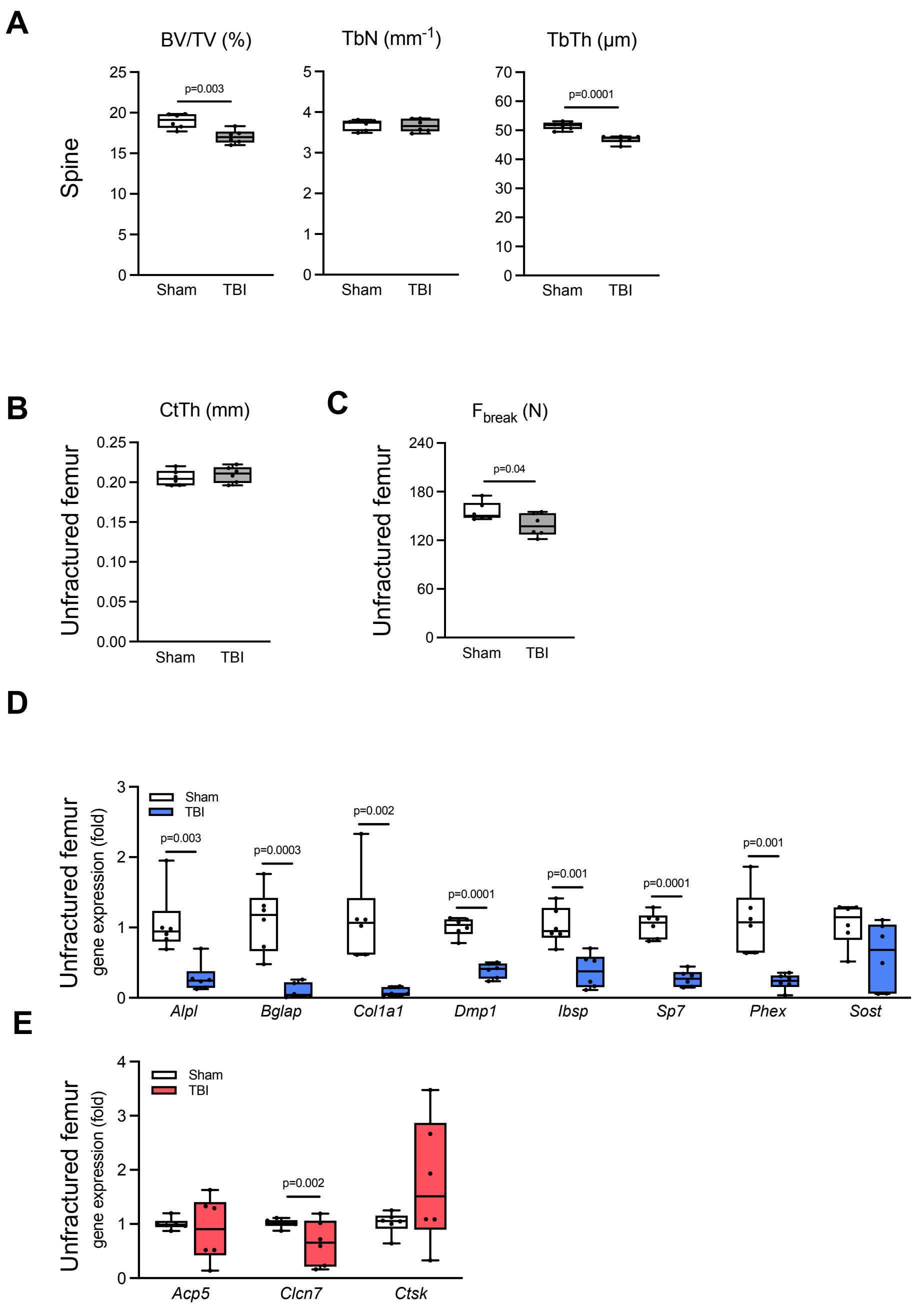


**Supplemental Figure 1. TBI causes systemic bone loss in the unfractured skeleton.** (**A**) Radiological quantification (µCT scanning) of bone volume per tissue volume (BV/TV), trabecular numbers (TbN), and trabecular thickness (TbTh) in the spine (L3 and L4) of 12-week-old female WT mice 14 days after TBI (FVB/129 genetic background). (**B**) Radiological quantification (µCT scanning) of cortical thickness in the femur midshaft of the same mice. (**C**) Biomechanical testing of unfractured femora derived from the same mice. $F_{break}$ = force to bone failure. (**D**) Expression of the indicated osteoblast marker genes in the unfractured femur of female mice (aged 12 weeks with FVB/129 genetic background) 3 days after TBI. *Alpl*, *alkaline phosphatase*; *Bglap*, *bone gamma-carboxyglutamate protein*; *Col1a1*, *collagen type I alpha 1*; *Dmp1*, *dentin matrix acidic phosphoprotein 1*; *Ibsp*, *bone sialoprotein*; *Sp7*, *osterix*; *Phex*, *phosphate regulating endopeptidase x-linked*; *Sost*, *sclerostin*. (**E**) Expression of the indicated osteoclast marker genes in the unfractured femur of the same mice. *Acp5*, *tartrate-resistant acid phosphatase type 5*; *Clcn7*, *chloride voltage-gated channel 7*; *Ctsk*, *cathepsin k*. Data are graphed in boxplots with median and 25$^{th}$ and 75$^{th}$ quantiles. Whiskers indicate upper and lower extremes, respectively. For all experiments, n=6 mice per group. For (**A**)-(**C**), two-tailed Student's t-test. For (**D**), (**E**) two-way ANOVA followed by Tukey's post hoc test.

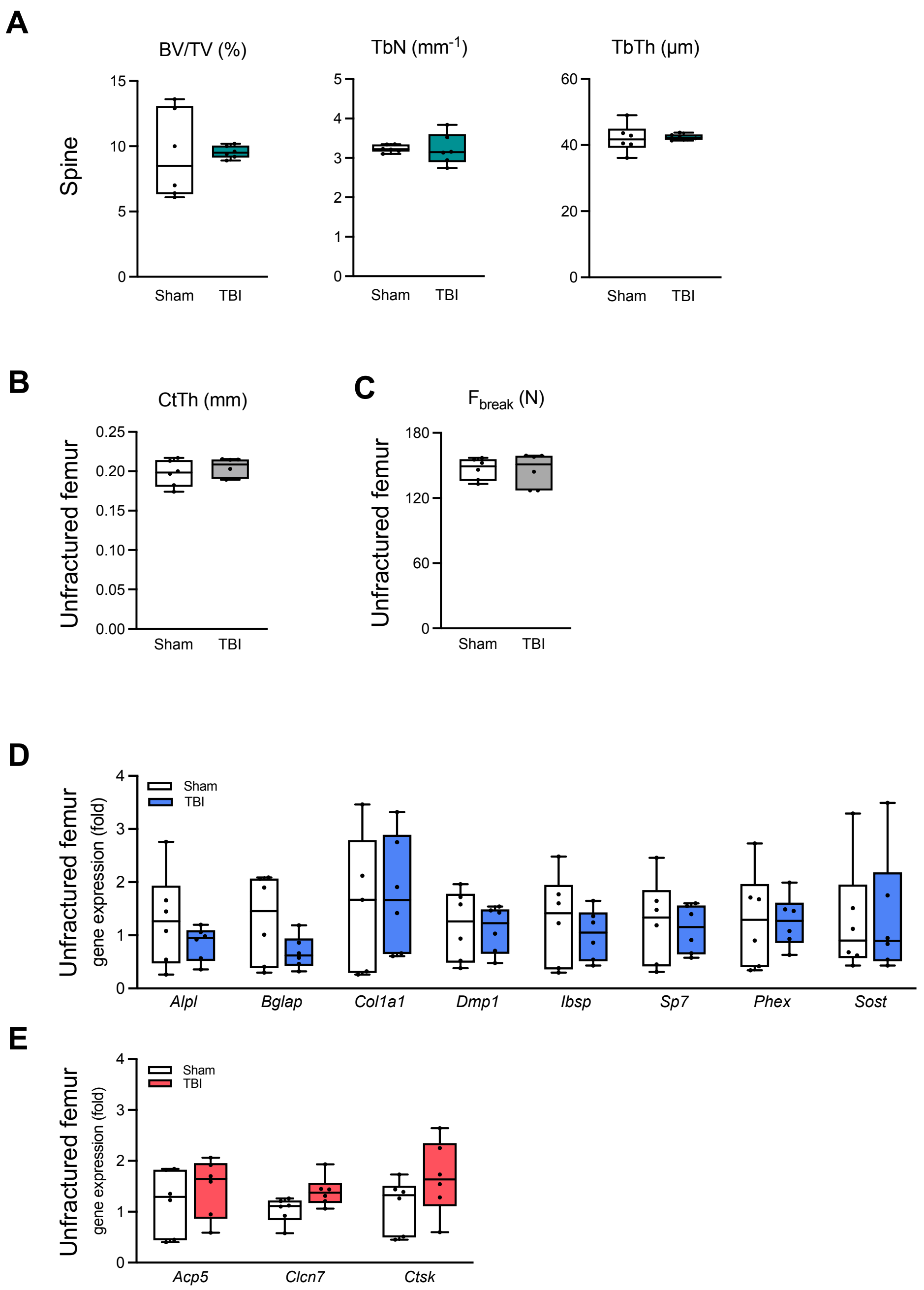


**Supplemental Figure 2. Adrb2-deficiency prevents the skeletal effects of TBI on bone remodeling and regeneration**. (**A**) Radiological quantification (µCT scanning) of bone volume per tissue volume (BV/TV), trabecular numbers (TbN), and trabecular thickness (TbTh) in the distal femur of 20-week-old female Adrb2-deficient mice 14 days after TBI (n=6 female mice per group, FVB/129 genetic background) (**B**) Radiological quantification (µCT scanning) of cortical thickness of the femur shaft in the same mice. (**C**) Biomechanical testing of unfractured femora of the same mice. $F_{break}$ = force to bone failure. (**D**) Expression of the indicated osteoblast marker genes in the unfractured femur of female mice (aged 12 weeks with FVB/129 genetic background) 3 days after TBI. *Alpl*, *alkaline phosphatase*; *Bglap*, *bone gamma-carboxyglutamate protein*; *Col1a1*, *collagen type I alpha 1*; *Dmp1*, *dentin matrix acidic phosphoprotein 1*; *Ibsp*, *bone sialoprotein*; *Sp7*, *osterix*; *Phex*, *phosphate regulating endopeptidase x-linked*; *Sost*, *sclerostin*. (**E**) Expression of the indicated osteoclast marker genes in the unfractured femur of the same mice. *Acp5*, *tartrate-resistant acid phosphatase type 5*; *Clcn7*, *chloride voltage-gated channel 7*; *Ctsk*, *cathepsin k*. Data are graphed in boxplots with median and $25^{th}$ and $75^{th}$ quantiles. Whiskers indicate upper and lower extremes, respectively. For all experiments, n=6 mice per group. For (**A**)-(**C**), two-tailed Student's t-test. For (**D**), (**E**) two-way ANOVA followed by Tukey's post hoc test.

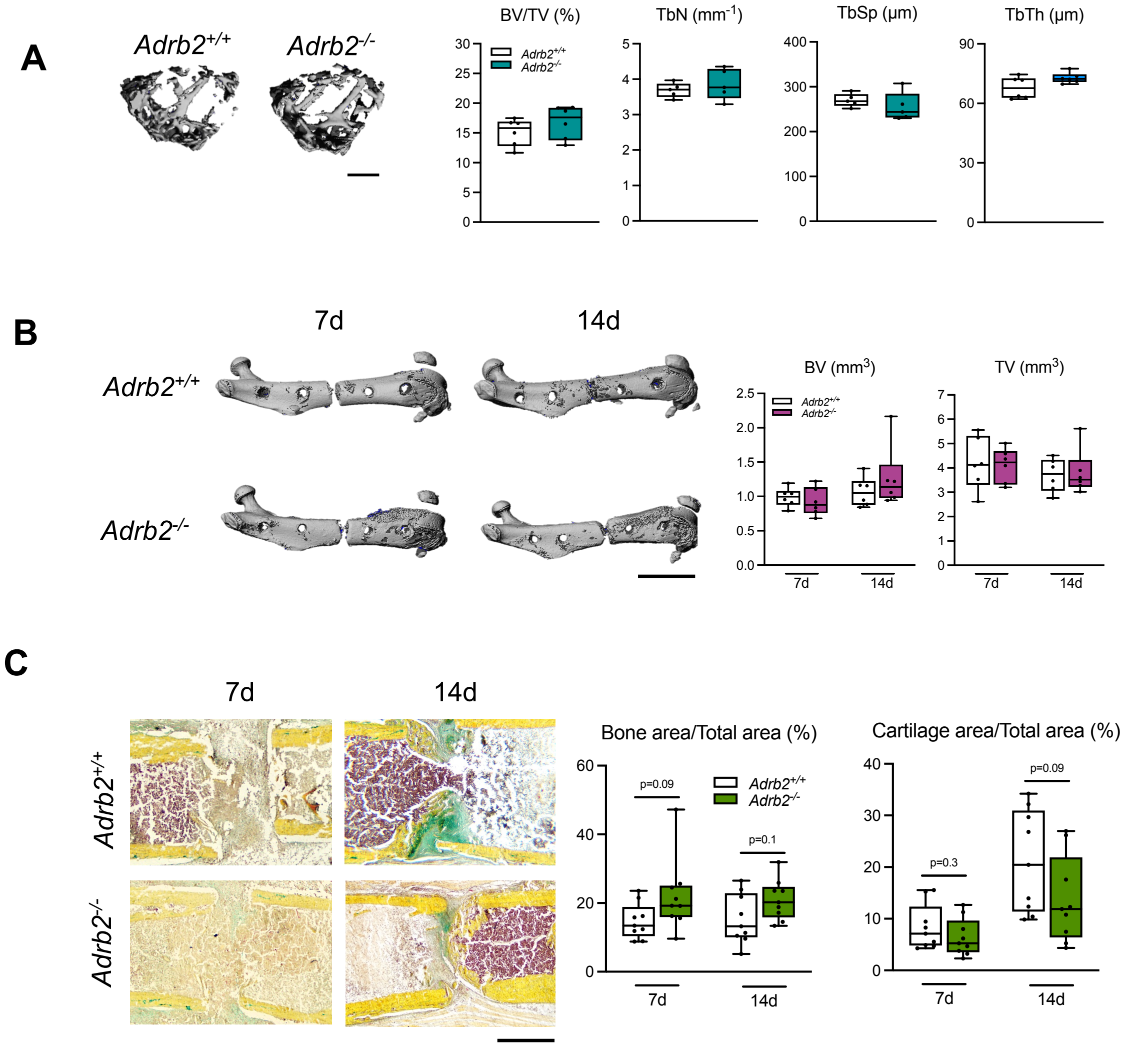


**Supplemental Figure 3. Insignificant role of Adrb2 in bone remodeling and fracture healing in young mice.** (**A**) Representative µCT images of the distal femora in male 12-week-old male mice (FVB/129 genetic background) of the indicated genotypes and radiological quantification of bone volume per tissue volume (BV/TV), trabecular numbers (TbN), trabecular separation (TbSp), and trabecular thickness (TbTh). Scale bar=200 µm. (**B**) Representative µCT images of the fractured femora in 12-week-old male mice (FVB/129 genetic background) of the indicated genotypes 7 and 14 days following fracture and quantification of callus bone volume (BV) and tissue volume (TV). Scale bar=5 mm. (**C**) Representative callus sections (Movat Pentachrome staining) in the same mice and quantification of bone area/total area and cartilage area/total area at the indicated time points. Scale bar=400 µm. Data are graphed in boxplots with median and 25$^{th}$ and 75$^{th}$ quantiles. Whiskers indicate upper and lower extremes, respectively. For all experiments, n=6-9 mice per group as indicated. For (**A**), two-tailed Student's t-test. For (**B**), (**C**) two-way ANOVA followed by Tukey's post hoc test.

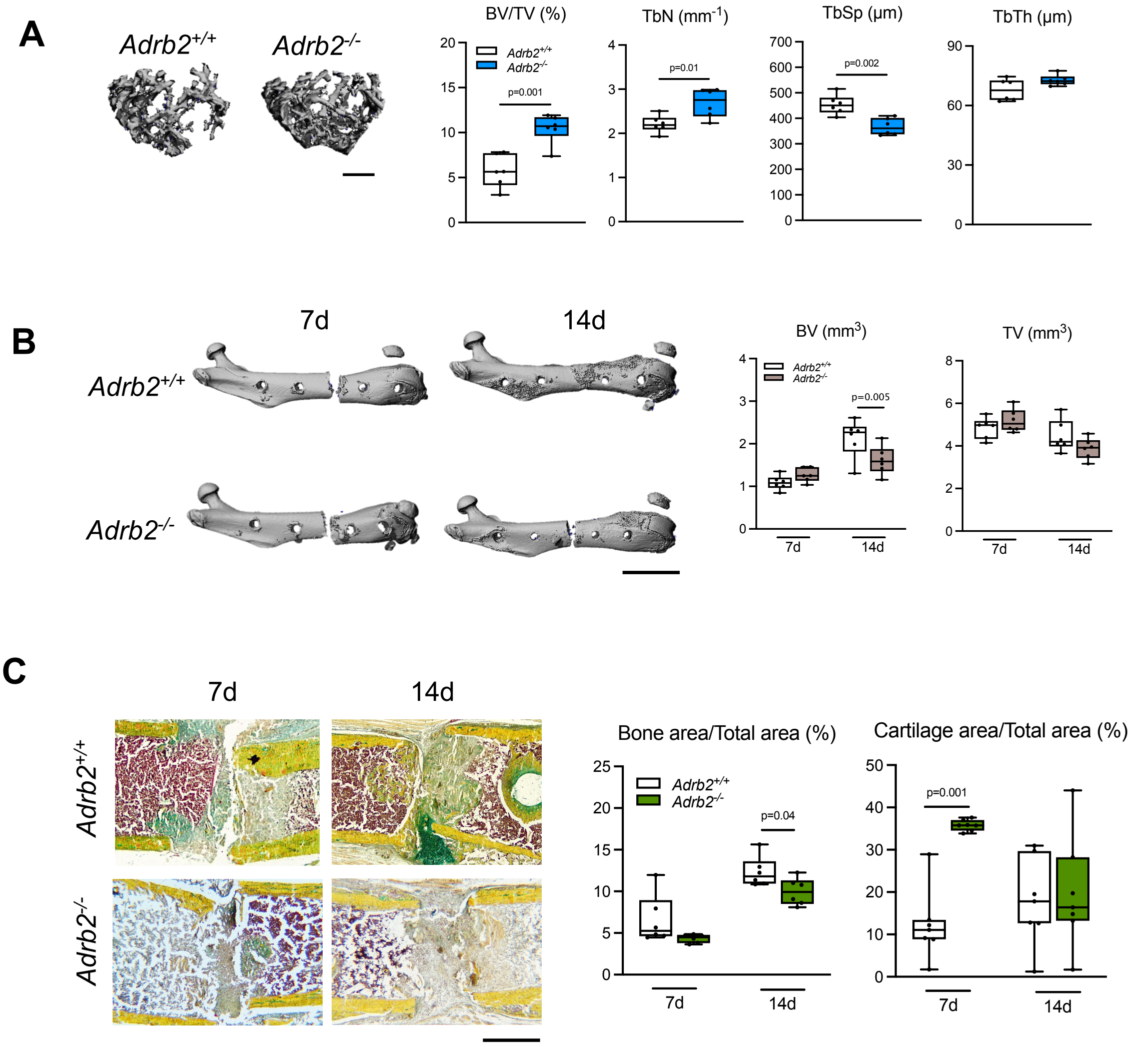


**Supplemental Figure 4. Age-dependent role of Adrb2 in fracture healing.** (**A**) Representative µCT images of the distal femora in 30-week-old male mice (FVB/129 genetic background) of the indicated genotypes and radiological quantification of bone volume per tissue volume (BV/TV), trabecular numbers (TbN), trabecular separation (TbSp), and trabecular thickness (TbTh). Scale bar=200 µm. (**B**) Representative µCT images of the fractured femora in 12-week-old male mice of the indicated genotypes 7 and 14 days following fracture and quantification of callus bone volume (BV) and tissue volume (TV). Scale bar=5 mm. (**C**) Representative callus sections (Movat Pentachrome staining) in the same mice and quantification of bone area/total area and cartilage area/total area at the indicated time points. Scale bar=400 µm. Data are graphed in boxplots with median and 25th and 75th quantiles. Whiskers indicate upper and lower extremes, respectively. For all experiments, n=6 mice per group. For (**A**), two-tailed Student's t-test. For (**B**), (**C**) two-way ANOVA followed by Tukey's post hoc test.

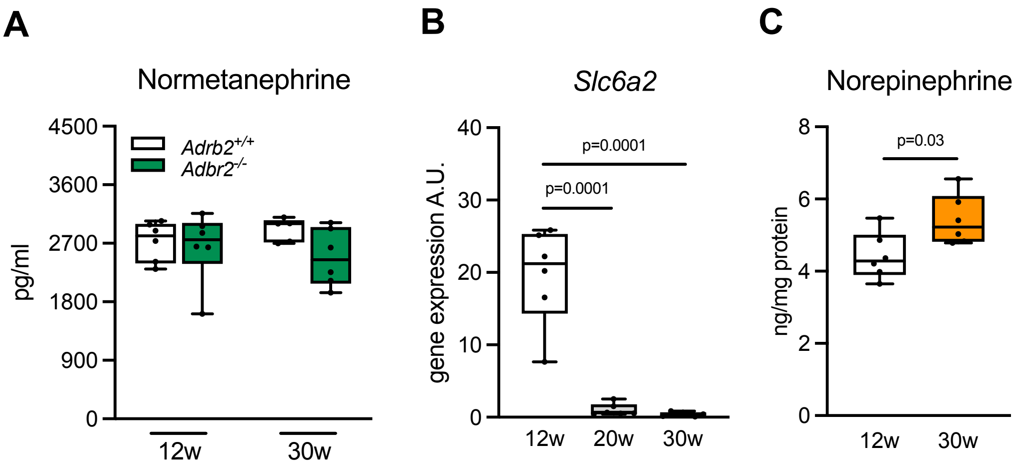


**Supplemental Figure 5. Indices of increased sympathetic tone in the femora of aged mice.** (**A**) Normetanephrine levels in serum of young (12-week-old) and aged (30-week-old) male WT mice measured by ELISA. (**B**) Expression of the NE transporter *Slc6a2* (*solute carrier family 6 member 2*) in flushed femora of the same mice measured by qRT-PCR. (**C**) NE content in the flushed femora of the same mice normalized to protein concentrations. Data are graphed in boxplots with median and 25th and 75th quantiles. Whiskers indicate upper and lower extremes, respectively. For all experiments, n=6 mice (FVB/129 genetic background) per group. For (**A**), (**B**) two-way ANOVA followed by Tukey's post hoc test. For (**C**), two-tailed Student's t-test.

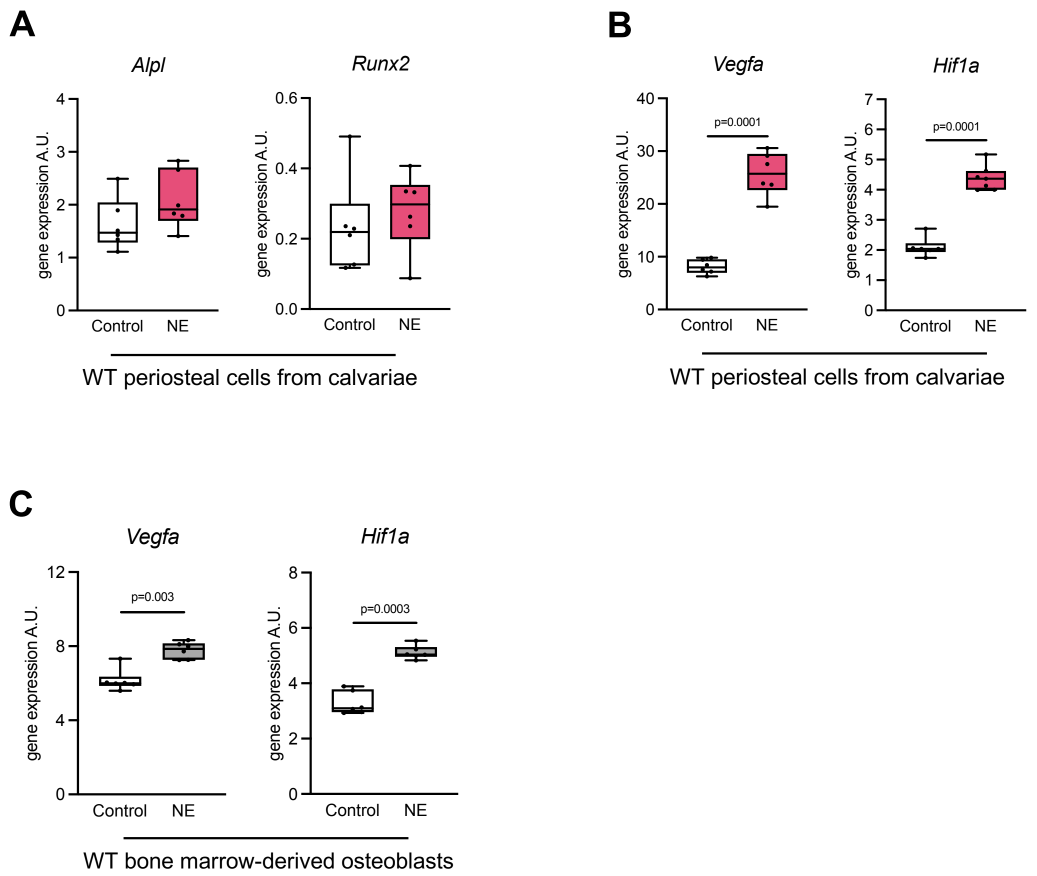


**Supplemental Figure 6. Effect of NE on gene expression in periosteal cells and osteoblasts.** (**A**) Expression of the indicated osteoblast marker genes in WT periosteal cells at day 15 of differentiation stimulated with/without NE for 6 hours. Cells were derived from calvarial bones of both male and female mice with a C57Bl/6J genetic background (12-18 weeks of age). (**B**) Expression of the indicated angiogenic marker genes in the same cells. (**C**) Expression of the indicated angiogenic marker genes in WT bone marrow-derived osteoblasts at day 10 of differentiation stimulated with/without NE for 6 hours. Cells were derived from femoral bone marrow of both male and female mice with FVB/129 genetic background (12-18 weeks old). Data are graphed in boxplots with median and 25$^{th}$ and 75$^{th}$ quantiles. Whiskers indicate upper and lower extremes, respectively. For all experiments, n=6 independent cell cultures per group. Two-tailed Student's t-test.

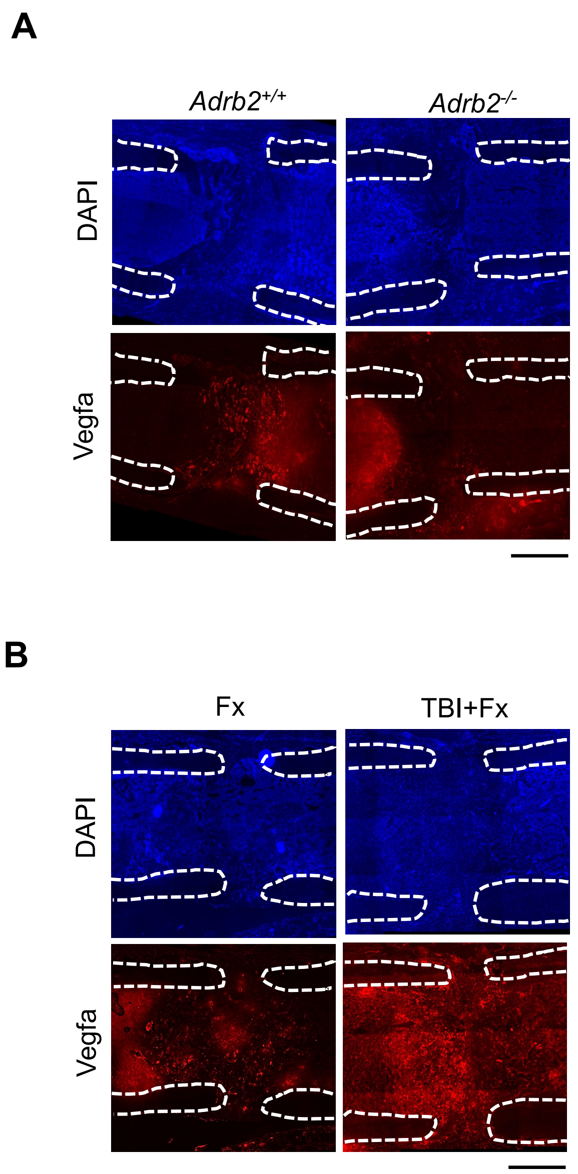


**Supplemental Figure 7. Unmerged images of Vegfa-specific immunofluorescence in the fracture callus.** (**A**) Representative immunofluorescent images using an Vegfa-specific antibody and DAPI stainings in the fracture callus of 30-week-old male mice of the indicated genotypes 7 days after osteotomy. Scale bar=300 µm. (**B**) Representative immunofluorescent images using an Vegfa-specific antibody in the fracture callus of 12-week-old female WT mice 7 days after osteotomy with/without TBI. Scale bars=300 µm.

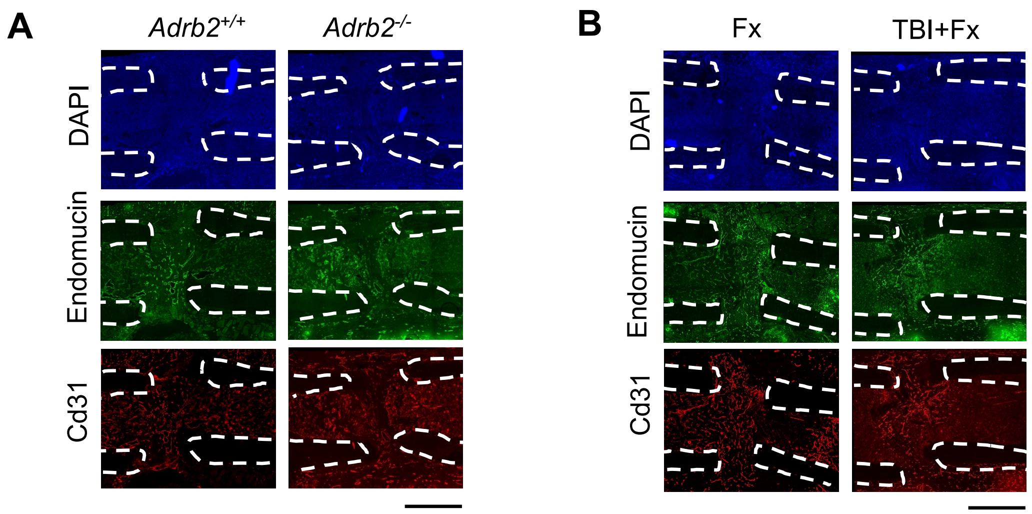


**Supplemental Figure 8. Unmerged images of Cd31+Endomucin co-immunofluorescence in the fracture callus.** (**A**) Representative images of unmerged immunofluorescent co-stainings using Cd31- and Endomucin-specific antibodies in the fracture callus of 30-week-old male mice of the indicated genotypes 7 and 14 days after osteotomy. (**B**) Representative images of unmerged immunofluorescent co-stainings using Cd31- and Endomucin-specific antibodies in the fracture callus of 12-week-old female WT mice 14 days after osteotomy with/without TBI. Scale bars=400 µm.

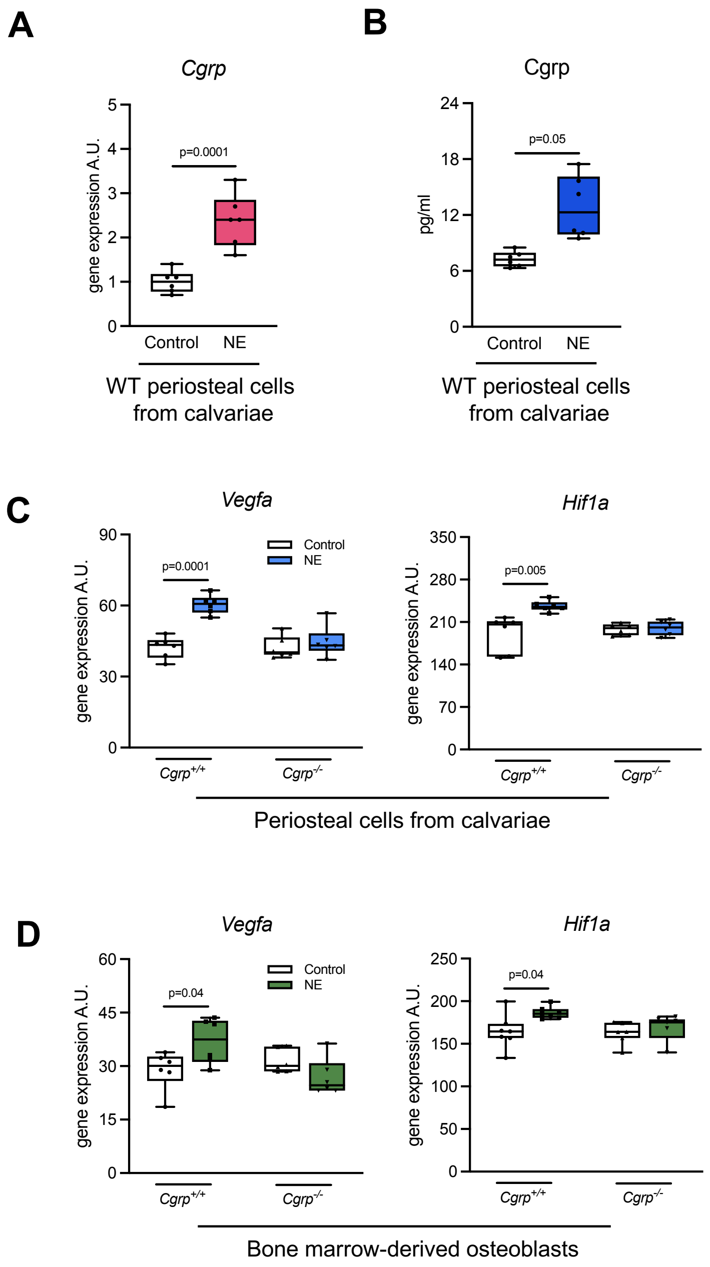


**Supplemental Figure 9. aCgrp-dependent induction of Vegfa by NE in periosteal cells and osteoblasts.** (**A**) Expression of *aCgrp* in WT periosteal cells after stimulation with NE for 6 hours and (**B**) Cgrp protein concentrations in the supernatant of the same cultures measured by ELISA. Cells were derived from calvarial bones of both male and female WT mice with FVB/129 genetic background (12-18 weeks of age). (**C**) Expression of the indicated genes in WT and aCgrp-deficient periosteal cells stimulated with NE for 6 hours. Cells were derived from calvariae of both male and female aCgrp-deficient mice with C57Bl/6J genetic background (12-18 weeks of age). (**D**) Expression of the indicated genes in WT and aCgrp-deficient bone marrow-derived osteoblasts stimulated with NE for 6 hours. Cells were derived from femoral bone marrow of both male and female aCgrp-deficient mice with C57Bl/6J genetic background (12-18 weeks of age). For (**A**)-(**D**), n=6 independent cultures per group. For (**A**), (**B**), two-tailed Student's t-test. For (**C**), (**D**), two-way ANOVA followed by Tukey's post hoc test.

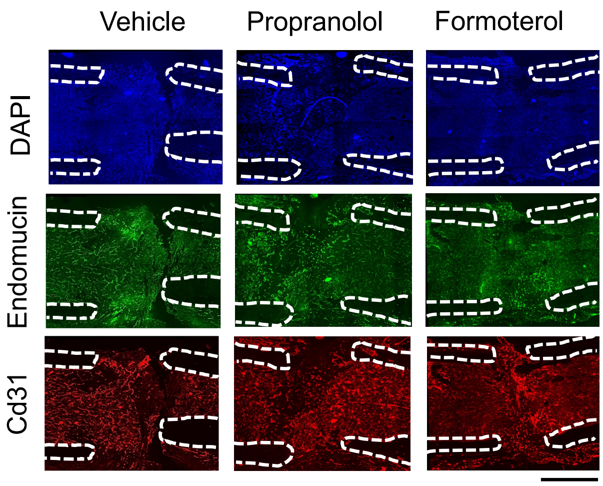


**Supplemental Figure 10. Unmerged images of Cd31+Endomucin co-immunofluorescence in the fracture callus of mice treated with Adrb2 antagonist and agonist.** Representative images of unmerged immunofluorescent co-stainings using Cd31- and Endomucin-specific antibodies in the fracture callus of 30-week-old male mice with the indicated treatments 14 days after osteotomy. Scale bar=400 µm.

| | **NE** (mean ± SD) | **No NE** (mean ± SD | **p** |
|---|---|---|---|
| n | 37 | 37 | - |
| Age | 39.45 ± 17.66 | 45.68 ± 19.61 | 0.097 |
| ISS | 17.89 ± 10.22 | 6.62 ± 4.05 | <0.001 |
| Cumulative hours NE iv | 51.61 ± 80.13 | - | - |

**Supplemental Table 1. Baseline characteristics of the study cohorts with long bone fractures.** ISS=injury severity score. Comparisons were made using two-tailed Student's t-test.
